# NADH regulation shapes competing acetate flux responses in *Escherichia coli*: kinetic modelling of the tricarboxylic acid cycle and acetate production

**DOI:** 10.64898/2026.11.20.768084

**Authors:** Jiarui Li, Mark B. Flegg, Vijay Rajagopal, Christopher Drovandi, Michael McDonald, Jennifer A. Flegg, Michael Pan

## Abstract

Overflow metabolism occurs when cells divert carbon towards fermentative products despite oxygen availability. In rapidly growing *Escherichia coli*, it is primarily characterised by acetate secretion. An elevated NADH/NAD^+^ ratio has been proposed to contribute to this behaviour, but the mechanisms linking redox imbalance to acetate production remain unclear. We developed a kinetic model linking pyruvate metabolism, the tricarboxylic acid (TCA) cycle, and acetate formation to examine how cofactor availability and NADH-mediated enzyme regulation influence carbon partitioning. The model combines thermodynamically consistent reversible rate laws with NADH inhibition of pyruvate dehydrogenase (PDH) and citrate synthase, and was calibrated using Bayesian inference to account for parameter uncertainty. We examined steady-state responses to changes in the target NADH/NAD^+^ ratio and the total NAD(H) pool size. With both inhibitory interactions included, lowering the target redox ratio increased acetate-producing flux, contrary to the experimentally observed reduction in acetate secretion. Substantially attenuating the effective NADH response of PDH recovered the experimental direction of this response. Additionally, to reproduce the observed reduction in acetate secretion following NAD(H)-pool expansion, a weak but non-zero PDH response was required. These findings suggest that cofactor-dependent kinetics and NADH-mediated regulation can jointly contribute to carbon redistribution between the TCA cycle and acetate production. Within this restricted model, whether a more reduced redox state promotes acetate-producing flux depends critically on the effective NADH sensitivity of PDH.

## 1 Introduction

Overflow metabolism, the secretion of incompletely oxidised carbon products despite the availability of oxygen, remains one of the most debated phenomena in physiology [1, 2]. It is observed across diverse biological systems, including bacteria, yeasts, and proliferating mammalian cells, including cancer cells [1]. In *Escherichia coli*, this phenomenon manifests as acetate overflow, where carbon flux is diverted from complete respiratory oxidation toward acetate secretion during rapid growth, even though complete respiratory oxidation provides a higher adenosine triphosphate (ATP) yield per glucose [2, 3]. Despite its ubiquity, the mechanistic basis of overflow metabolism remains unclear, and a number of hypotheses have been proposed to explain why cells exhibit this apparently inefficient phenotype [1, 4]. These explanations include proteome allocation constraints [3], membrane space limitations [5], and redox balancing mechanisms [6]. No single mechanism appears sufficient to explain overflow across all physiological conditions, motivating the computational investigation of individual proposed mechanisms.

Acetate is produced mainly from acetyl-CoA via the phosphate acetyltransferase (Pta)–acetate kinase (AckA) pathway [7]. Acetate overflow in *E. coli* can therefore be viewed, from one perspective, as carbon partitioning at the acetyl-CoA branch point between oxidation through the tricarboxylic acid (TCA) cycle and acetate production through Pta–AckA. This framing is particularly relevant to redox-based explanations because routing carbon through the TCA cycle is tightly coupled to the generation of reducing equivalents (NADH, ubiquinol), whereas diversion of acetyl-CoA toward acetate reduces the generation of additional reducing equivalents downstream of acetyl-CoA and may therefore lower the burden on respiratory NADH oxidation [8].

Redox balance offers one potential explanation for overflow metabolism in *E. coli*. During the oxidation of carbon sources such as glucose, the nicotinamide adenine dinucleotide couple (NAD^+^/NADH) carries the resulting reducing equivalents, with NAD^+^ serving as an electron acceptor [9]. The subsequent re-oxidation of NADH to NAD^+^ is essential for maintaining redox balance and sustaining continuous cellular growth. Under aerobic conditions, oxygen serves as the terminal electron acceptor, allowing NADH to be oxidised through the respiratory chain. In contrast, under anaerobic conditions where alternative electron acceptors are absent, NAD^+^ regeneration is achieved through fermentation pathways that reduce metabolic intermediates using NADH in *E. coli* [9]. Therefore, changes in NADH availability are expected to influence the distribution of metabolic fluxes, even under aerobic conditions. Moreover, Vemuri et al. reported a sharp increase in the intracellular NADH/NAD^+^ ratio near the onset of overflow metabolism [10]. This observation motivated the hypothesis that an elevated redox ratio can contribute to the regulation of overflow metabolism, rather than merely reflect saturation of the electron transport system [10]. A range of mechanisms has been proposed to explain this link. One is redox-responsive transcriptional regulation through ArcAB, whose activity is linked to the respiratory redox state and which represses TCA cycle and respiratory genes under reducing conditions [10]. Another is that overflow arises when the demand for NAD^+^ regeneration outpaces the demand for ATP production, such that the coupled respiratory chain cannot regenerate NAD^+^ quickly enough and fermentation becomes favourable even in the presence of oxygen [6]. Although this framework was developed in eukaryotic systems, it suggests a broader perspective: redox requirements may influence carbon allocation even when this reduces ATP yield or carbon efficiency.

Despite this range of proposed mechanisms, it remains unclear what generates a change in whole-cell redox state, and how a given change in redox state may translate into acetate-producing flux. Notably, acetate formation has been found to persist even after ArcA-mediated transcriptional repression was removed. This suggests that overflow metabolism could arise through an alternative mechanism, such as direct inhibition of citrate synthase (CS) by NADH [10]. In other words, ArcA-mediated transcriptional regulation may not be required for the immediate flux response, which may instead be driven by local biochemical regulation, reactant availability, and thermodynamic constraints. The TCA cycle and its entry reactions are particularly relevant in this context because NADH directly inhibits pyruvate dehydrogenase (PDH) and CS, which control acetyl-CoA production and its entry into the cycle, respectively [11, 12]. Moreover, because NADH and NAD^+^ participate directly in several dehydrogenase reactions, changes in the NADH/NAD^+^ ratio alter the corresponding thermodynamic driving forces [13]. This extensive coupling to the cellular redox state motivates our focus on the TCA cycle.

Systems biology models have a long history in the investigation of metabolic function [14, 15].Stoichiometric models capture the structural connectivity of metabolic networks and can predict plausible flux distributions. However, they do not by themselves explain how metabolite availability, enzyme regulation, and redox balance interact dynamically to produce acetate overflow [16, 17]. This limitation motivates the use of mechanistic dynamic frameworks that integrate these processes within the underlying metabolic network.

Numerous studies have developed kinetic and related mechanistic representations of the TCA cycle and broader central carbon metabolism under different modelling assumptions, ranging from small pathway models to large networks calibrated for specific growth conditions [16, 18–23]. More directly relevant to acetate overflow, Millard et al. [2] developed a six-species, six-reaction coarse-grained model linking glucose uptake and lumped glycolytic production of acetyl-CoA to a lumped TCA cycle flux coupled to biomass formation and an explicit reversible Pta–AckA pathway with acetate exchange. Acetate-dependent inhibition of the lumped glycolytic and TCA cycle capacities was required to reproduce steady-state and dynamic acetate-flux behaviours; however, because individual TCA cycle reactions and intracellular NAD(H) pools were not represented, the model could not resolve which molecular or redox-sensitive interactions mediated these responses [2]. These models have provided valuable insights into local regulatory interactions and pathway-level dynamics. Some studies, including Millard et al., have propagated measurement uncertainty through repeated model fitting [2]. Given the sparsity of kinetic data, parameters may remain weakly constrained or strongly correlated. Even when marginal confidence intervals are reported, it remains important to assess structural and practical identifiability, joint parameter dependencies, and the robustness of mechanistic conclusions across independent perturbations. These issues are particularly important when kinetic models are extrapolated beyond the conditions used for calibration. It therefore remains unclear whether NADH-dependent kinetics linking pyruvate metabolism, the TCA cycle, and acetate production can reproduce the observed direction of acetate-flux responses to independent experimental perturbations, and whether this conclusion remains robust to the residual parameter uncertainty after calibration.

In the past decade, there has been growing interest in combining Bayesian inference with kinetic metabolic modelling to address these limitations. A seminal example is the framework of Saa and Nielsen, who constructed thermodynamically feasible kinetic ensembles and updated them with data using approximate Bayesian computation, thereby obtaining approximate posterior distributions over parameters and predictions rather than a single calibrated model [24]. More recent work has extended this idea to larger networks and richer data types, including genome-scale linlog-kinetic models inferred from multi-omics datasets, computational platforms such as GRASP for constructing thermodynamically feasible kinetic ensembles, and Bayesian ordinary differential equation (ODE) frameworks for tracer-based non-steady-state metabolomics data [25–27]. Together, these studies show that Bayesian approaches can provide mechanistically interpretable kinetic models together with principled uncertainty quantification.

In this work, we use a mathematical model to investigate whether direct NADH-dependent regulation within the acetate fermentation and TCA cycle pathways is sufficient to reproduce the observed directions of acetate-flux responses to experimentally imposed redox perturbations. By treating upstream carbon input as a boundary process and representing respiratory NADH oxidation through a lumped reaction, we isolate how a prescribed change in the NADH/NAD+ state is translated into carbon partitioning between TCA-associated oxidation and acetate production through the Pta–AckA pathway. We formulate this module using thermodynamically consistent convenience kinetics, including NADH inhibition of PDH and CS, and calibrate it to steady-state metabolite and flux data using Bayesian inference. We then simulate complementary perturbations of the NADH/NAD^+^ ratio and the total NAD(H) pool across posterior parameter samples. Within the model, the experimentally observed directions of the acetate-flux responses to both perturbations can be reconciled only when PDH retains a weak but non-zero effective sensitivity to NADH; stronger PDH inhibition primarily suppresses acetyl-CoA throughput, whereas NADH regulation of CS more directly redistributes flux between the TCA cycle and acetate production.

## 2 Methods

An overview of the modelling and inference workflow is shown in Figure 1. Section 2.1 describes the model structure and kinetic rate laws. Section 2.2 details the parameterisation procedure, including parameter balancing, a Bayesian method for integrating kinetic and thermodynamic information to obtain a thermodynamically consistent kinetic parameter set. Section 2.3 assesses structural and practical identifiability and parameter recovery using synthetic data. Section 2.4 describes calibration of the model to experimental steady-state data using Bayesian inference and evaluation using posterior predictive checks. Section 2.5 presents the global sensitivity analysis used to quantify how parameter variation influences model outputs.

**Figure 1:**
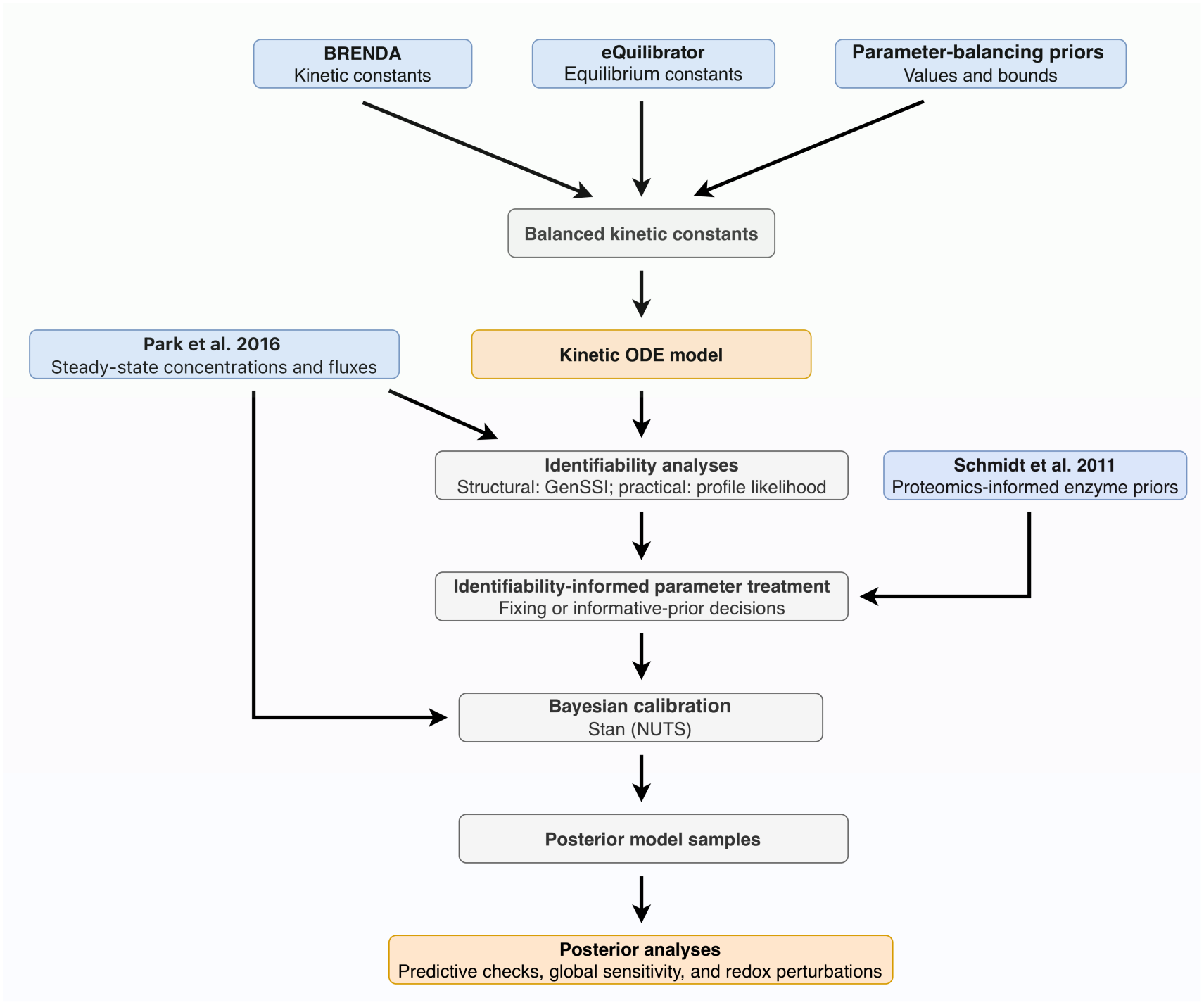
Overview of the modelling and inference workflow. Kinetic parameters are assembled from literature data and thermodynamic constraints using parameter balancing. The resulting model is analysed for structural identifiability, followed by practical identifiability and parameter estimation using synthetic and then experimental data. Bayesian inference and posterior predictive checks are used to assess model performance. Abbreviations: ODE, ordinary differential equation; NUTS, No-U-Turn Sampler; GenSSI, Generating Series for testing Structural Identifiability.

### 2.1 Model formulation

We developed a reduced dynamic model linking pyruvate metabolism, the TCA cycle, and acetate production in *E. coli*, described by a system of ODEs for the concentrations of 15 intracellular metabolites in the TCA cycle and acetate overflow modules. The reaction topology was adapted from EColiCore2 [28] and included pyruvate kinase, pyruvate dehydrogenase, citrate synthase, two sequential aconitase steps, isocitrate dehydrogenase, *α*-ketoglutarate dehydrogenase, succinyl-CoA synthetase, succinate dehydrogenase, fumarase, malate dehydrogenase, phosphate acetyltransferase, and acetate kinase (Figure 2).

**Figure 2:**
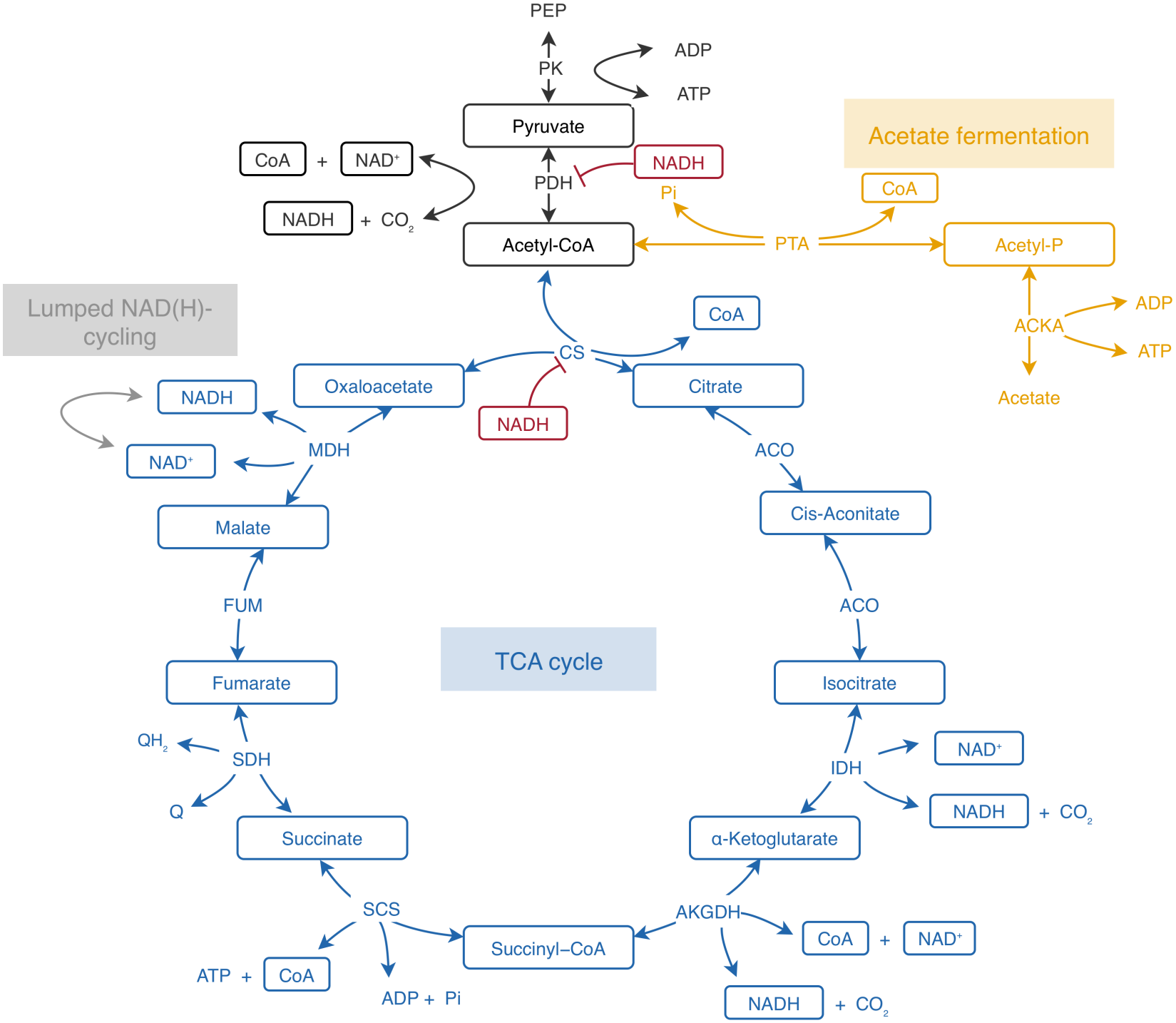
Schematic of the reduced kinetic model linking pyruvate metabolism, the TCA cycle and acetate production in *E. coli*. The model comprises pyruvate entry (black), the TCA cycle (blue), the acetate-producing Pta–AckA pathway (orange), and a lumped bidirectional NAD(H)-cycling process (grey). Metabolite species enclosed in boxes are dynamic state variables, whereas unboxed species are fixed boundary variables. Enzymecatalysed reactions are shown as arrows, whereas NADH-mediated inhibition of PDH and CS is shown by blunt-ended lines (red). PEP, phosphoenolpyruvate; CoA, coenzyme A; Q, ubiquinone; QH_2_, ubiquinol; ADP, adenosine diphosphate; Pi, inorganic phosphate; CO_2_, carbon dioxide; Acetyl-P, acetyl phosphate; PK, pyruvate kinase; PDH, pyruvate dehydrogenase; CS, citrate synthase; ACO, aconitase; IDH, isocitrate dehydrogenase; AKGDH, α-ketoglutarate dehydrogenase; SCS, succinyl-CoA synthetase; SDH, succinate dehydrogenase; FUM, fumarase; MDH, malate dehydrogenase; PTA, phosphate acetyl-transferase; ACKA, acetate kinase.

The model was designed to isolate the local response of the pyruvate–acetyl-CoA branch, where NADH-sensitive reactions can alter carbon partitioning between TCA-associated oxidation and acetate production through the Pta–AckA pathway. Upstream glycolytic supply was represented through pyruvate entry from boundary metabolites, while whole-cell NADH oxidation via other pathways and NAD^+^ regeneration were represented by lumped phenomenological NAD(H)-cycling terms.

Let 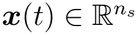 denote the vector of dynamic intracellular metabolite concentrations (mM), where *n*_*s*_ = 15 is the number of species, and let ***p*** denote the complete set of model parameters and constants. These include Michaelis constants *K*_*m*_ (mM), catalytic turnover rates *k*_cat_ (s^*−*1^), inhibition constants *K*_*i*_ (mM), effective enzyme abundances *E* (mM), redox-related parameters, and boundary (fixed) metabolite concentrations (mM). The model is written as

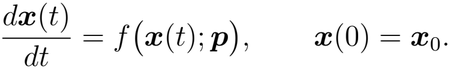

Specific subsets of ***p*** were varied or estimated in the identifiability, calibration, and sensitivity analyses, as described below; all remaining quantities were held fixed. Expanding the ODE system explicitly, the mass-balance equations are given by

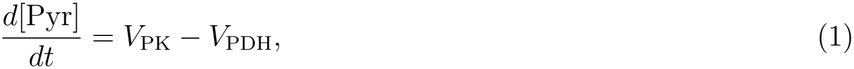

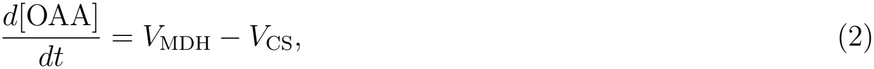

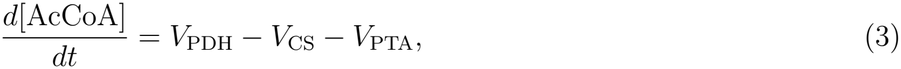

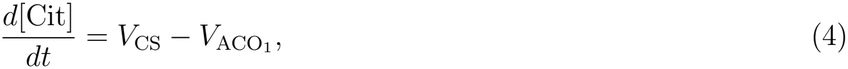

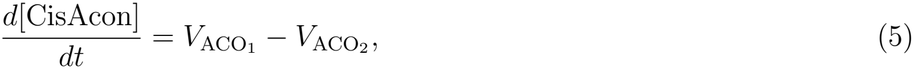

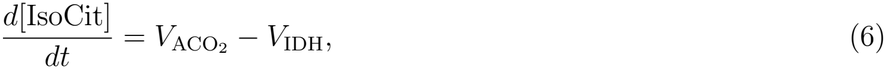

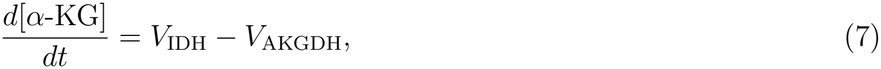

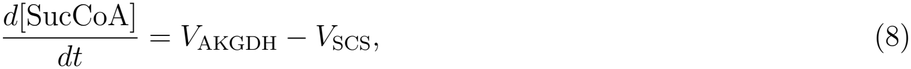

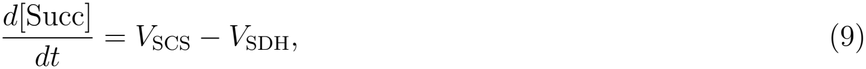

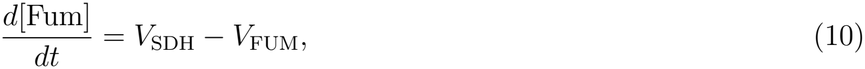

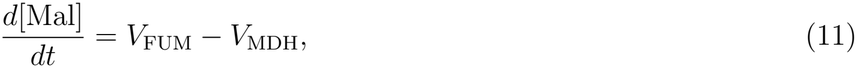

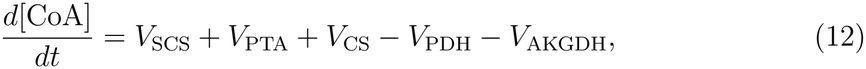

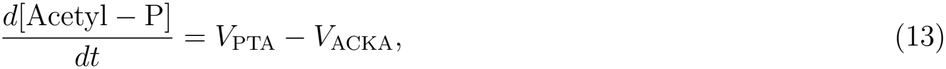

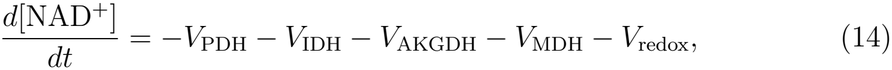

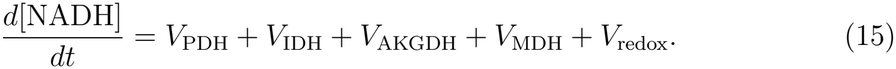

Here, *V*_PK_, *V*_PDH_, … denote the corresponding reaction fluxes. The enzyme-catalysed reaction fluxes were represented using convenience kinetics [29], which generalises reversible Michaelis–Menten kinetics to enzyme-catalysed reactions with arbitrary stoichiometries and multiple substrates and products. For the reactions considered here, whose stoichiometric coefficients are unity, the rate of a generic reaction is written as

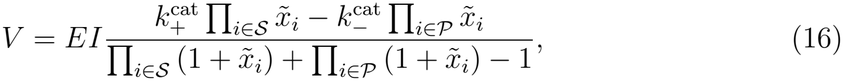

where *S* and *P* denote the substrate and product sets, respectively, *E* is the effective enzyme abundance, 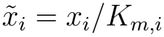 is the normalised concentration of metabolite *i*, and 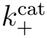 and 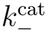 are the forward and reverse catalytic turnover numbers. The factor *I* accounts for inhibition, under the assumption that it is independent of enzyme catalysis. For an inhibitor *j*, we have

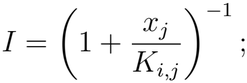

otherwise, *I* = 1. The specific factors representing NADH inhibition of PDH and CS are denoted by *I*_PDH_ and *I*_CS_, respectively.

The lumped phenomenological flux *V*_redox_, which is not an enzyme-catalysed reaction flux, is defined separately in Equation (61) in Supplementary Section S1.3. Reaction-specific expressions for all enzyme-catalysed fluxes *V*_*i*_ are provided in the Supplementary Section S1.2. Boundary metabolite concentrations and enzyme abundances were held constant during each simulation. Growth-associated dilution was omitted from the metabolite concentration balances under the assumption that intracellular metabolite turnover was fast relative to dilution by cellular growth.

### 2.2 Parameterisation and calibration data

Kinetic parameters were estimated using parameter balancing, which integrates experimental measurements and log-normal priors within a Bayesian framework to accommodate missing data and generate thermodynamically consistent parameter sets [30, 31]. The framework enforces Haldane relationships linking reversible kinetic parameters to reaction equilibrium constants, and Wegscheider conditions ensuring consistency of equilibrium constants across stoichiometrically closed reaction cycles. Measurements were compiled from the literature and the BRENDA database [32], while equilibrium constants were obtained from eQuilibrator [33]. Details of parameter balancing are provided in the Supplementary Section S2.1.

From the complete model parameter vector ***p***, we defined an initial candidate vector ***θ***_cand_ containing the quantities whose values were not independently known. These comprised effective enzyme-abundance parameters, redox-cycling parameters, and unmeasured boundary-metabolite concentrations.

Structural identifiability analysis was first used to obtain a reduced parameter vector ***θ***_str_. Practical identifiability analysis was then used to determine the final subset ***θ***_cal_ retained for Bayesian calibration. All remaining components of ***p*** were fixed at their parameter-balanced, literature-based, or independently specified values. These included the Michaelis constants *K*_*m*_, catalytic turnover rates *k*_cat_, and inhibition constants *K*_*i*_. The source kinetic and thermodynamic data and the parameter-balanced values are reported in Supplementary Table S1. The candidate, successive reduction process and final retained parameter sets are described in Table 1 in Supplementary Section S2.2.

**Table 1:** Successive reduction of the candidate parameter set based on structural identifiability analysis. Structural identifiability was assessed under ideal continuous-time, noise-free observations with known initial conditions.

| Step | Parameter set | No. of free parameters | Result | Action taken |
|---|---|---|---|---|
| 1 | Full candidate parameter set | 17 | The complete parameter vector was not structurally identifiable. Most parameters were globally identifiable; *E*<sub>SDH</sub>, *E*<sub>ACKA</sub>, and *x*<sub>Acetate</sub> were locally identifiable; and *x*<sub>*Q*</sub> and *x*<sub>*QH*<sub>2</sub></sub> were non-identifiable. | Fixed *x*<sub>*Q*</sub> and *x*<sub>*QH*<sub>2</sub></sub> at their assumed physiological values. |
| 2 | First reduced parameter set | 15 | No parameters remained non-identifiable. All parameters were globally identifiable except *x*<sub>Acetate</sub> and *E*<sub>ACKA</sub>, which remained locally identifiable. | Fixed *x*<sub>Acetate</sub> at the baseline boundary value and retained *E*<sub>ACKA</sub> as an unknown parameter. |
| 3 | Final reduced parameter set | 14 | All remaining parameters were structurally globally identifiable. | Retained the final parameter set for practical identifiability analysis. |

Experimental calibration data were obtained from Park et al. for *E. coli* K-12 NCM3722 grown at 37^*°*^C in Gutnick minimal medium containing D-glucose. Absolute intracellular metabolite concentrations and steady-state fluxes were estimated in exponentially growing filter cultures [34]. Fluxes reported in mM min^*−*1^, normalised to aqueous cell volume, were converted to mM s^*−*1^ by division by 60 before comparison with model predictions. The experimental metabolite concentrations and pathway fluxes, together with their reported units and available lower and upper bounds, are provided in Supplementary Table S2.

The stoichiometric matrix gives rise to three independent conserved totals, identified from its left null space [35]: the total coenzyme A pool, the total TCA-intermediate pool, and the total NAD(H) pool:

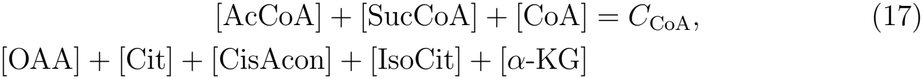

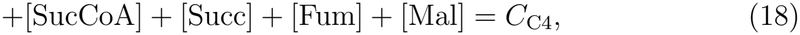

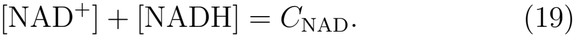

The values of these conserved totals were determined by the initial conditions and remained invariant along simulated model trajectories. These equalities are structural conservation relations rather than separately imposed state constraints.

### 2.3 Identifiability analysis

Kinetic metabolic models often contain many weakly constrained parameters and can exhibit sloppy parameter sensitivities, posing substantial challenges for parameter inference and model validation [36]. Recent work has emphasised that meaningful parameter inference in systems biology requires explicit assessment of structural and practical identifiability to determine whether candidate parameters are learnable in principle and from finite, noisy data, respectively [37]. Accordingly, we first assessed structural identifiability under idealised observation conditions and then evaluated practical identifiability using profile likelihood. These analyses were used to define the parameter subset retained for subsequent Bayesian calibration.

#### 2.3.1 Structural identifiability

Before parameter estimation, we assessed structural identifiability to determine which candidate parameters could, in principle, be uniquely recovered from idealised, continuous, noise-free observations of the chosen outputs under the specified initial conditions and fixed model quantities [36].

Consider an ODE system

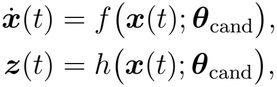

where ***θ***_cand_ denotes the candidate unknown parameter vector, and ***z***(*t*) denotes the vector of model outputs used for the identifiability analysis. All fixed model quantities are suppressed from the notation, and the initial conditions are treated as known. The model is globally structurally identifiable if the mapping

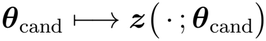

is injective over the admissible parameter space, such that no two distinct candidate parameter vectors produce identical output trajectories. It is locally structurally identifiable if, for a given admissible choice of parameter values, no distinct parameter vector within a neighbourhood of that choice produces the same output trajectory [36].

Structural identifiability was analysed using the differential–algebraic framework implemented in the GenSSI toolbox for Matlab [38]. GenSSI constructs an output-generating series from successive Lie derivatives,

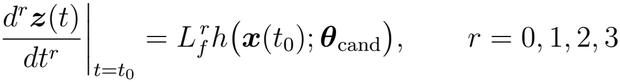

which encodes the information available from ideal observations.

The candidate parameter vector ***θ***_cand_ comprised effective enzyme-abundance parameters, redox-cycling parameters, and boundary-metabolite concentrations that were treated as unknown; measured boundary concentrations, initial conditions, and all other fixed model quantities were treated as known. The complete candidate parameter and output vectors supplied to GenSSI are given in Supplementary Section 2.2, and the successive parameter-set reductions are summarised in Table 1. Fixed boundary concentrations and conserved pool totals are listed in Supplementary Table S4.

GenSSI translates the output-generating series into algebraic constraints and examines their solvability to classify parameters as globally identifiable, locally identifiable, or non-identifiable [38]. The resulting classifications were used to obtain the structurally reduced parameter vector ***θ***_str_ before practical identifiability analysis and Bayesian calibration. Because structurally non-identifiable quantities cannot be uniquely estimated from the available outputs, one parameter within each non-identifiable combination was fixed at a nominal value to remove the corresponding redundancy. The parameters fixed in this way and their nominal values are listed in Supplementary Table S1.

#### 2.3.2 Practical identifiability and profile likelihood

Structural identifiability is a necessary condition for unique parameter estimation from ideal noise-free observations, but it does not guarantee that parameters can be estimated precisely from finite, noisy, or sparse data. Even structurally identifiable parameters may therefore remain practically non-identifiable under a particular observation scheme [39]. We assessed practical identifiability under the steady-state observation scheme using profile-likelihood analysis applied to the experimental metabolite-concentration and reaction-flux data used for Bayesian calibration. A separate synthetic-data analysis was performed to assess parameter recovery under controlled conditions and is reported in Supplementary Section S2.3.1.

Let ***θ***_str_ denote the vector of retained parameters, and let *J* (***θ***_str_) denote the negative log-likelihood used as the optimisation objective. For each parameter of interest *θ*_str,*i*_, we partition the parameter vector into *θ*_str,*i*_ and the remaining parameters ***θ***_str,*−i*_. With this partition, we write *J* (***θ***_str_) = *J* (*θ*_str,*i*_, ***θ***_str,*−i*_). The profile negative log-likelihood is then defined as

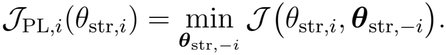

The minimisation is performed over the remaining parameters within their prescribed bounds, while *θ*_str,*i*_ is held fixed. Numerically, each parameter was fixed successively over a predefined range while the remaining parameters were re-optimised by minimising the negative log-likelihood, equivalently by maximising the profile log-likelihood [40].

For presentation, the profile was expressed as a normalised profile log-likelihood difference:

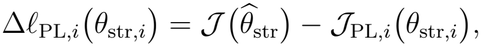

where 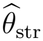 denotes the maximum-likelihood estimate. By construction, Δℓ_PL,*i*_ = 0 at the maximum-likelihood estimate and decreases as *θ*_str,*i*_ is constrained away from its optimum. An approximate 95% profile-likelihood confidence set was constructed using the standard asymptotic likelihood-ratio criterion [40]:

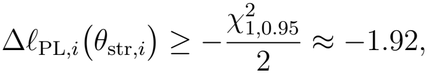

where 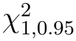 denotes the 95th percentile of the chi-square distribution with one degree of freedom.

A profile crossing this threshold on both sides of its maximum was classified as two-sided practically recoverable under the experimental steady-state observation scheme. Profiles crossing the threshold on only one side were classified as weakly practically identifiable, whereas profiles that did not cross the threshold in either direction were classified as practically non-identifiable [39]. The analysis was used to identify parameters that were weakly or one-sidedly constrained by the experimental likelihood and for which posterior inference could therefore be sensitive to prior specification. These parameters received additional weakly informative prior regularisation, with prior locations and scales informed by available biochemical or physiological information. All parameters in ***θ***_str_ were retained in the final Bayesian calibration vector ***θ***_cal_.

### 2.4 Bayesian parameter estimation

Bayesian parameter inference was performed in `Stan` through `CmdStanR` using the No-U-Turn Sampler (NUTS) [41, 42]. For each proposed calibration parameter vector ***θ***_cal_, the complete model parameter vector ***p*** was assembled by combining ***θ***_cal_ with the fixed model parameters. The ODE system was integrated from the specified initial conditions to *t*_end_ = 86,400 s (24 h) using Stan’s backward differentiation formula (BDF) solver, with relative tolerance 10^*−*3^, absolute tolerance 10^*−*5^, and a maximum of 2 *×* 10^5^ integration steps. The terminal-time state

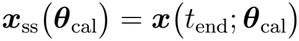

was used as the steady-state approximation, with fixed model parameters suppressed from the notation. The predicted metabolite concentrations corresponding to the calibration observations and the associated steady-state reaction fluxes are denoted by ***c***(***θ***_cal_) and ***v***(***θ***_cal_), respectively.

Metabolite concentrations and the fluxes considered here are positive-valued quantities subject to approximately multiplicative experimental uncertainty [43, 44]. Observations were therefore modelled on the log_10_ scale. A common log-scale standard deviation *σ*_*c*_ was used for concentration observations, whereas a separate standard deviation *σ*_*f*_ was used for flux observations:

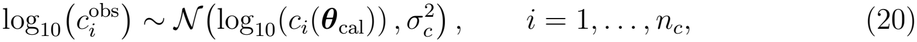

and

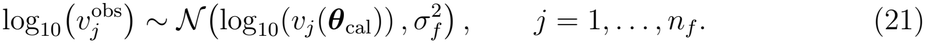

Here, *n*_*c*_ and *n*_*f*_ denote the numbers of concentration and flux observations, respectively. The error parameters *σ*_*c*_ and *σ*_*f*_ were estimated jointly with ***θ***_cal_. The prior treatment of the calibration parameters was informed by the profile-likelihood analysis. Parameters with two-sided profiles crossing the likelihood-ratio threshold were assigned broader priors, whereas parameters that were weakly or one-sidedly constrained received additional weakly informative regularisation based on available proteomic information reported by Schmidt et al. [45]. The complete prior specifications are provided in Supplementary Section S2.4.1. Four independent chains were run, each with 1,000 warmup iterations followed by 8,000 sampling iterations, yielding 32,000 post-warmup draws. Sampling was performed using `adapt_delta` = 0.9 and `max_treedepth` = 12. Convergence and sampling quality were assessed using rank-normalised 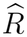 values, bulk and tail effective sample sizes [46], divergent transitions [47], and maximum tree-depth warnings [48]. Posterior draws were propagated through the kinetic model to quantify uncertainty in the predicted metabolite concentrations and reaction fluxes.

### 2.5 Global sensitivity analysis

Following Bayesian calibration, global sensitivity analysis was performed to quantify the effects of parameter variation on predicted steady-state metabolite concentrations and reaction fluxes. This analysis supported interpretation of model behaviour and did not alter the set of fixed or estimated parameters.

The set of quantities varied in the sensitivity analysis was not restricted to the parameters estimated during Bayesian calibration; it also included selected model quantities that were held fixed during calibration. They were collected into a factor vector ***ξ*** = (*ξ*_1_, … , *ξ*_*d*_), comprising 12 effective enzyme-abundance parameters, the NADH inhibition constants for PDH and CS, the total NAD(H) pool, the target redox parameter *α*, and the redox-cycling capacity *k*.

For sensitivity factors estimated during Bayesian calibration, baseline values were defined as the corresponding posterior medians; for factors held fixed during calibration, the values used in the calibrated model, obtained from the literature, were retained as baselines. For each factor *ξ*_*i*_ with baseline value *ξ*_*i*,0_, the sampling interval was [0.1*ξ*_*i*,0_, 10*ξ*_*i*,0_], with samples distributed uniformly on the log_10_ scale. Two *n × d* Sobol base matrices, **A** and **B**, were generated and used to construct the cross-sampled matrices 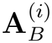 required by the Saltelli estimators.

For a scalar steady-state model output *Y* = *G*(***ξ***), the first-order Sobol index for factor *ξ*_*i*_ is

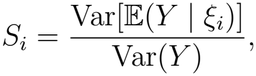

and the total-order index is

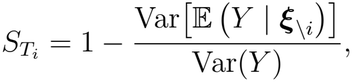

where ***ξ***_*\i*_ denotes all sensitivity factors except *ξ*_*i*_. The first-order index quantifies the proportion of variance in *Y* attributable to the main effect of *ξ*_*i*_, whereas the total-order index quantifies the overall contribution of *ξ*_*i*_, including its interactions with all other factors.

A total of *n*(*d* + 2) = 190,000 model evaluations were made for *n* = 10,000 base-sample rows and *d* = 17 sensitivity factors. The Sobol indices were computed separately for each steady-state metabolite concentration and reaction flux. Parameters with large first-order indices were interpreted as making substantial main-effect contributions to output variance, whereas differences between total-order and first-order indices indicated contributions from parameter interactions. Because Sobol indices are variance-based and unsigned, they do not indicate the direction of a factor’s effect on an output.

### 2.6 Redox perturbations and regulatory-response configurations

We considered two *in silico* redox perturbations. To represent changes associated with NADH oxidase overexpression, the target redox parameter *α* was varied relative to its sample-specific posterior value while the total NAD(H) pool was held fixed. To represent expansion of the intracellular NAD(H) pool, *C*_NAD_ was varied while *α* was held at its sample-specific posterior value. For each perturbation, the model was integrated to steady state at each imposed value.

Following the baseline-preserving approach of Tran et al. [49], we disabled a regulatory response while retaining its effect at the calibrated baseline state. For each posterior sample *s*, we first calculated the unperturbed steady state with both NADH-dependent inhibitory responses enabled and recorded the corresponding inhibition factors, 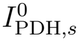 and 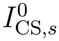 . During each redox perturbation, an enabled inhibition factor was evaluated using the current model-predicted NADH concentration, whereas a disabled factor was held at its sample-specific baseline value. Thus, disabling a regulatory response removed the model’s sensitivity to perturbation-induced changes in NADH while preserving its baseline inhibitory effect. Apart from the perturbed quantity, *α* or *C*_NAD_, all model quantities were held at their sample-specific posterior or fixed baseline values.

This procedure produced four regulatory-response configurations: both PDH and CS responses enabled, the PDH response enabled with the CS response held at baseline, the CS response enabled with the PDH response held at baseline, and both responses held at baseline.

## 3 Results

### 3.1 Bayesian-calibrated model reproduces observed steady-state concentrations and fluxes

We first used structural and practical identifiability analyses to determine the parameter set used for Bayesian calibration (overview in Figure 3). The initial candidate set comprised 17 unknown quantities, including effective enzyme-abundance parameters, redox-cycling parameters, and unmeasured boundary-metabolite concentrations. On the basis of the structural identifiability analysis, 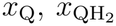 , and *x*_Acetate_ were fixed, leaving 14 globally structurally identifiable parameters for calibration.

**Figure 3:**
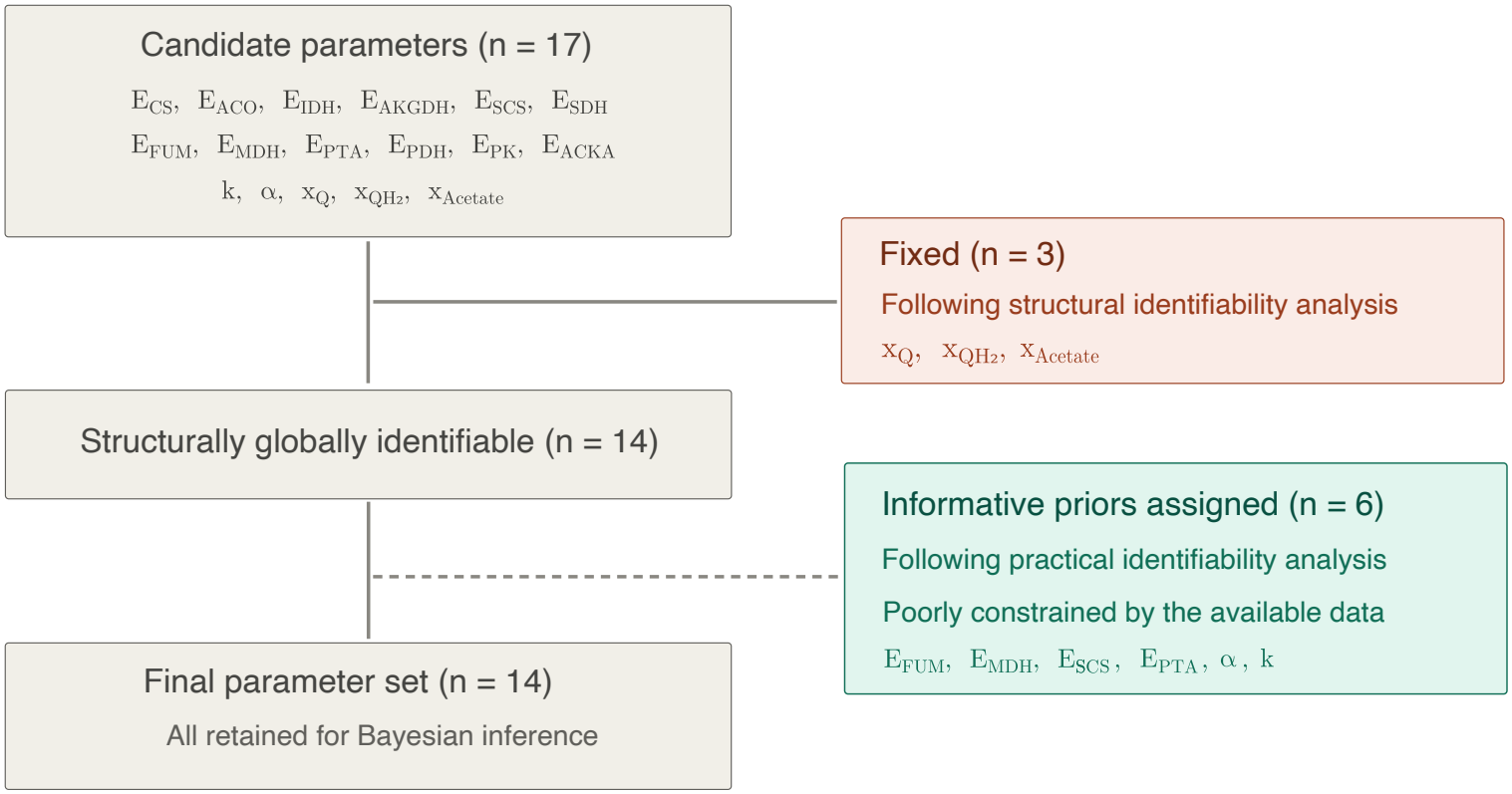
Reduction and regularisation of the parameter set used for Bayesian calibration. Structural identifiability analysis reduced the initial set of 17 candidate quantities to 14 globally structurally identifiable parameters. Practical identifiability analysis subsequently identified six poorly constrained parameters, which were retained for Bayesian inference with informative priors.

Profile-likelihood analysis of the experimental data did not lead to any further parameter removal (Figure S4). The profiles for *E*_MDH_, *k*, and *α* crossed the likelihood-ratio threshold on only one side, whereas those for *E*_SCS_, *E*_FUM_, and *E*_PTA_ did not cross the threshold in either direction. These parameters were retained in the Bayesian calibration, with more informative prior distributions used to regularise the estimation of weakly constrained parameters. The remaining eight parameters showed two-sided profiles and were assigned broader priors.

Having established the parameterisation, we calibrated the model against the steady-state metabolite concentration and flux measurements reported by Park et al. [34]. Posterior predictive checks were then used to assess agreement between the calibrated model and the observed data (Figure 4). The corresponding prior and posterior parameter distributions are shown in Figure 5.

**Figure 4:**
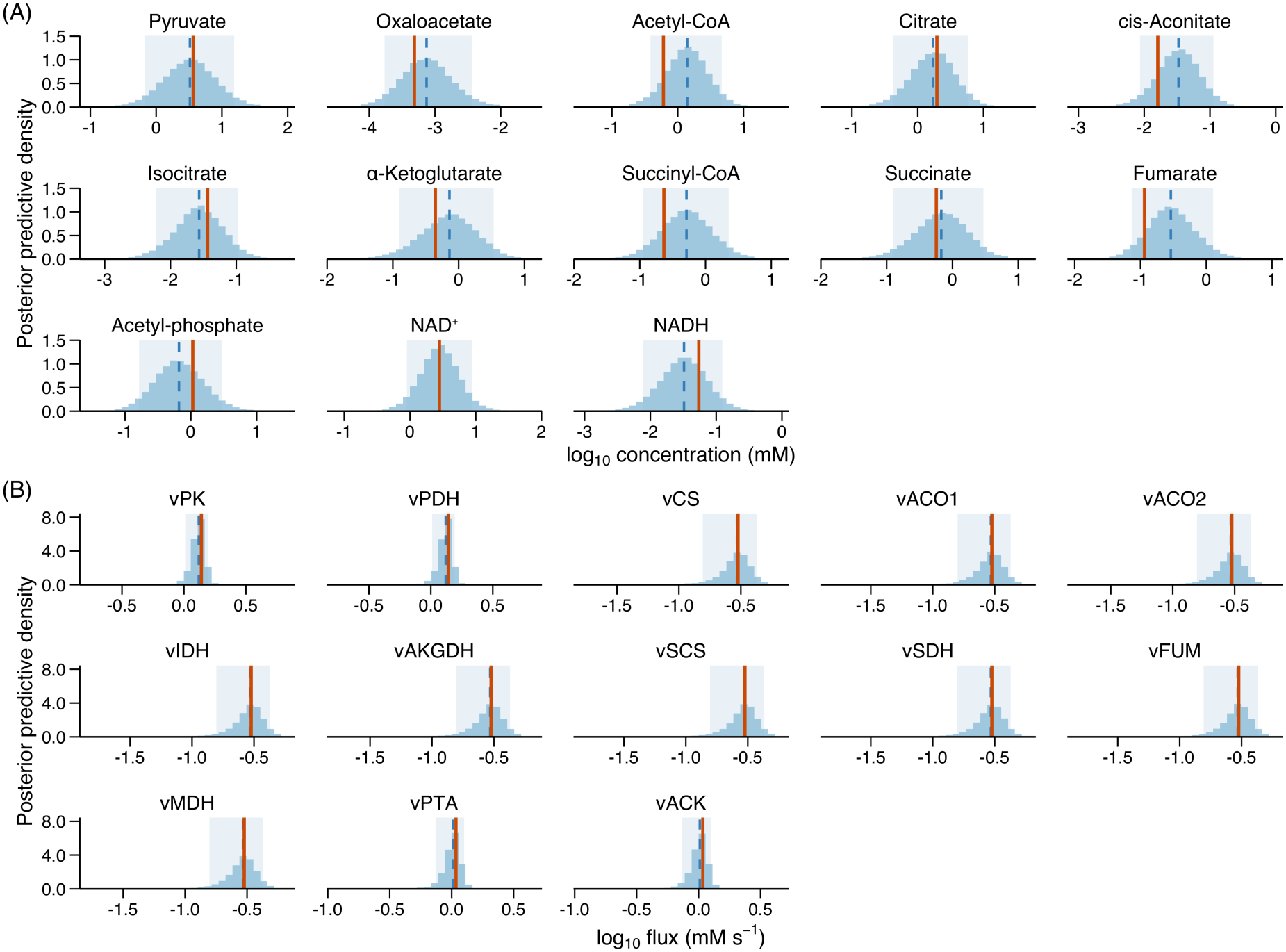
Posterior predictive checks for the steady-state TCA cycle–fermentation model fitted to experimental data. For each observed metabolite concentration and flux, the posterior predictive distribution is compared with the corresponding observed value. Shaded regions indicate the 90% posterior predictive intervals, red lines indicate the observed data, and blue dashed lines indicate the posterior predictive medians. Overall, the model reproduces the measured steady-state quantities with reasonable agreement.

**Figure 5:**
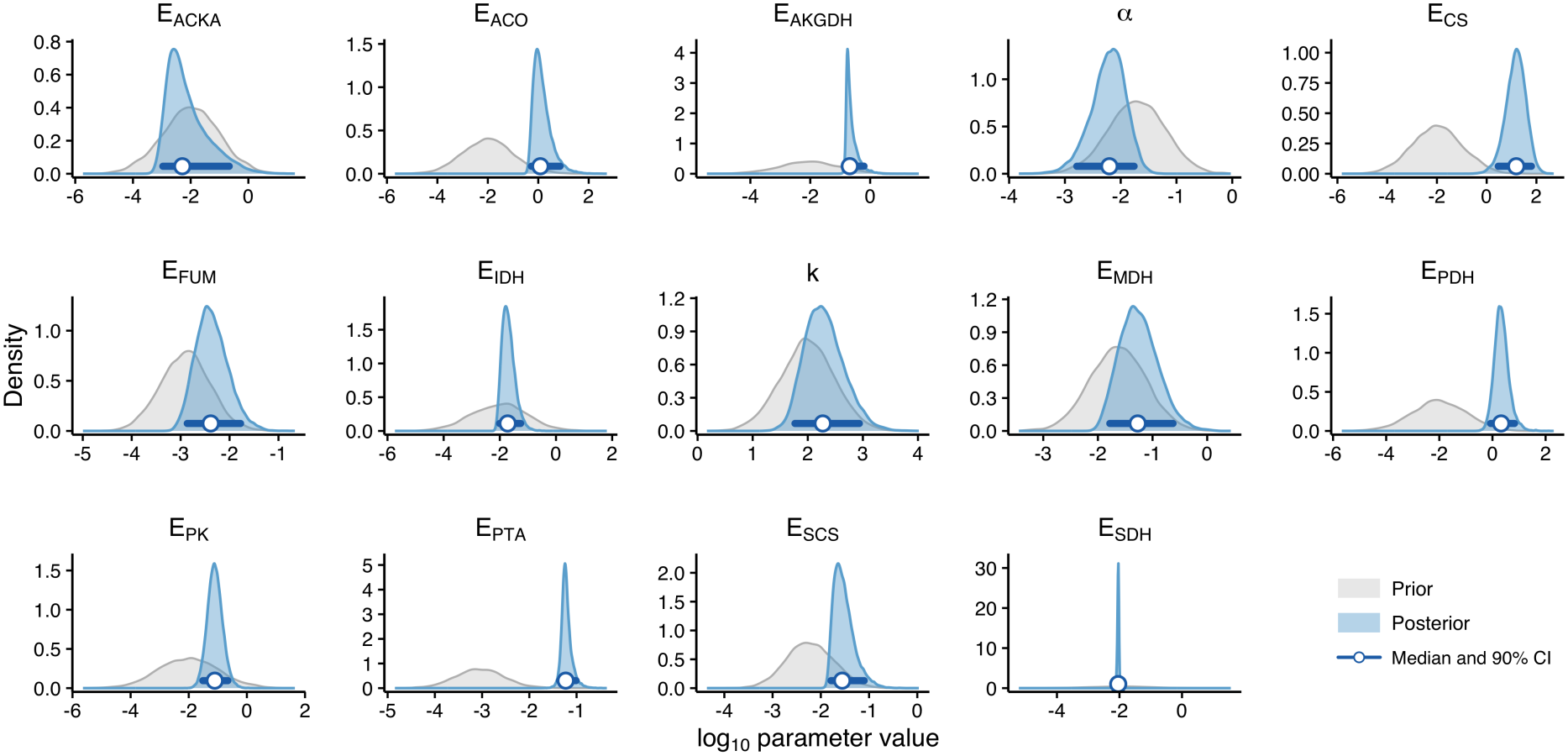
Prior and posterior distributions for the inferred model parameters in the steady-state TCA-fermentation model. Blue curves show the posterior distributions, grey curves show the prior distributions, and horizontal intervals with dots indicate the posterior 90% credible intervals and posterior medians, respectively.

Sampling diagnostics indicated satisfactory convergence and sampling efficiency across the four chains. All rank-normalised 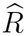 values were below 1.01, and the minimum bulk and tail effective sample sizes across the inferred parameters were 6,194 and 4,327, respectively. No divergent transitions occurred among the post-warm-up draws, and no transitions reached the maximum tree depth. Parameter-specific diagnostic summaries are provided in Supplementary Section S2.4.2.

In Figure 4, each panel compares the posterior predictive distribution for one measurement with its observed value. An observation falling within the shaded 90% posterior predictive interval indicates broad consistency between the fitted model and the corresponding measurement. The observed values for most metabolite concentrations and fluxes lie within their respective predictive intervals, although the observed fumarate concentration lies close to the lower edge of its interval. The predictive distributions within each measurement class have broadly comparable widths, although the flux predictions are generally more concentrated than the metabolite concentration predictions, indicating lower posterior predictive uncertainty for the fluxes. The similarity in spread within each class is partly due to the use of shared observation-noise parameters, *σ*_*c*_ for all metabolite concentrations and *σ*_*f*_ for all flux measurements, such that the spread of an individual posterior predictive distribution does not solely indicate how strongly the corresponding model prediction is constrained. Overall, the model provides an adequate description of the observed concentrations and fluxes of this experimental condition while yielding an ensemble of thermodynamically consistent and data-compatible baseline models for the subsequent perturbation analysis.

The prior-to-posterior comparison (Figure 5) shows posterior contraction for many of the calibrated parameters. However, concentrated posteriors are also observed for parameters whose experimental-data profile likelihoods are one-sided or do not cross the likelihood-ratio threshold (Figure S4). For these weakly constrained parameters, including *E*_SCS_, *E*_FUM_, *E*_PTA_, *E*_MDH_, *k*, and *α*, the more informative priors may contribute to posterior concentration. Their posterior widths should therefore not be interpreted as evidence of strong constraints from the experimental data alone.

### 3.2 Global sensitivity analysis reveals broad redox effects and pathway-specific flux sensitivities

Posterior predictive checks (Figure 4) showed that the calibrated model ensemble broadly reproduced the observed steady-state data. We next used variance-based global sensitivity analysis to determine how variation in effective enzyme capacities and redox-related parameters propagated to predicted steady-state metabolite concentrations and fluxes. The resulting total-order Sobol indices, which incorporate both main effects and parameter interactions, are shown in Figure 6.

**Figure 6:**
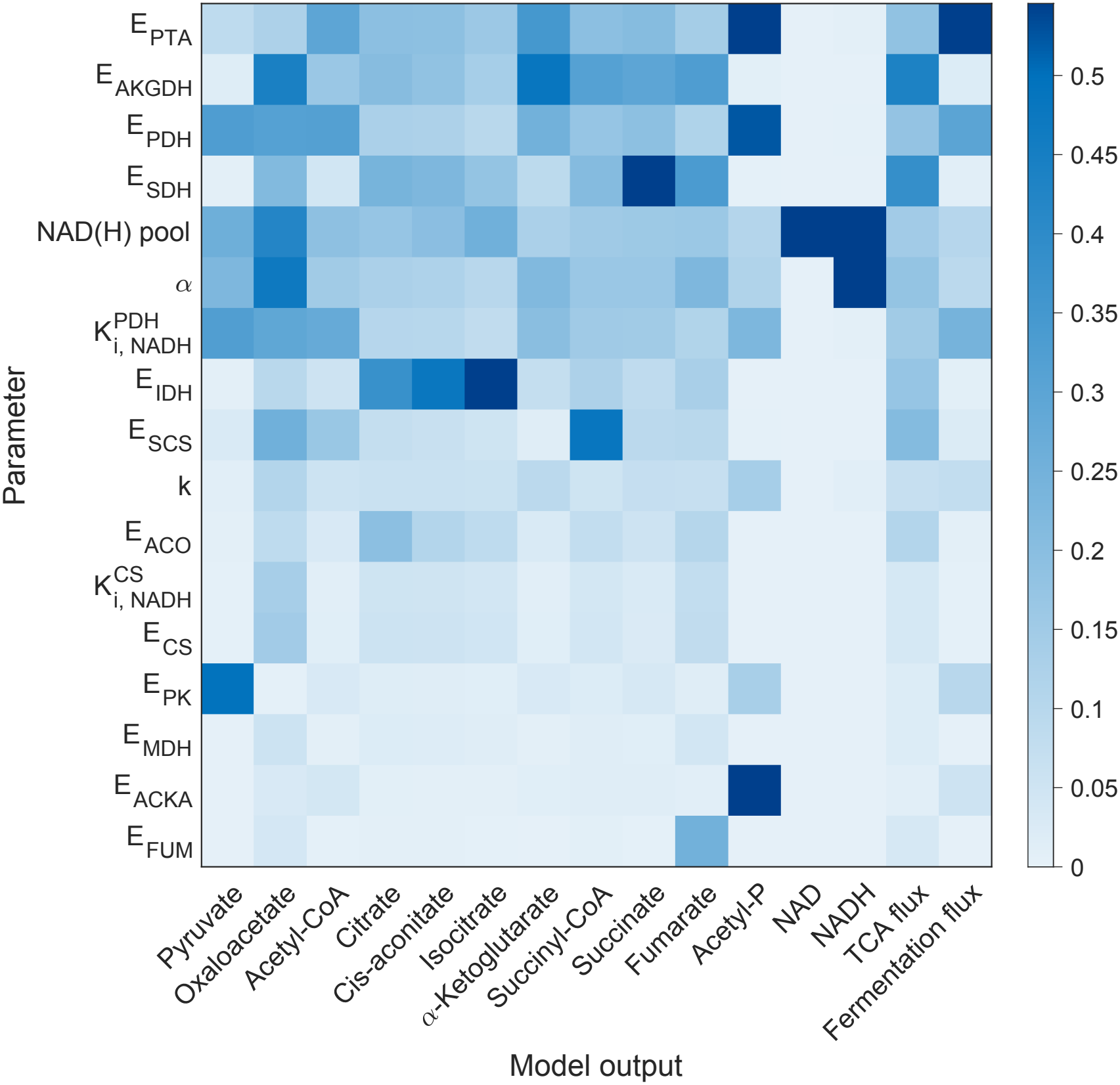
Global sensitivity of steady-state model predictions to effective enzyme capacities and redox-related parameters. Heatmap entries show the total-order Sobol index for each parameter–output pair, incorporating both the parameter’s main effect and all interactions involving that parameter. Parameters were varied independently over a 100-fold range, from 0.1 to 10 times their baseline values, using log-uniform sampling and 10,000 Sobol base samples. Darker shading indicates greater sensitivity.

The sensitivity analysis distinguished broad redox-mediated effects from pathway-specific sensitivity patterns. The total NAD(H) pool and the target redox parameter *α* showed the broadest influence across the model outputs. Among the effective enzyme-capacity parameters, *E*_AKGDH_ and *E*_SDH_ were among the strongest influences on TCA cycle flux. Fermentation flux was most sensitive to *E*_PTA_, consistent with the direct role of Pta at the entry to the acetate-producing branch. *E*_PDH_ and its NADH inhibition constant, 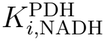 , also exerted substantial effects on fermentation flux, whereas the corresponding CS-related parameters had markedly weaker effects. Other enzyme-capacity parameters showed more localised sensitivity patterns, most notably *E*_IDH_ across citrate, cis-aconitate, and isocitrate, *E*_SCS_ for succinyl-CoA, *E*_FUM_ for fumarate, and *E*_ACKA_ for acetyl phosphate.

### 3.3 Effects of PDH and CS inhibition on responses to the target NADH/NAD^+^ ratio

The Sobol analysis indicated that PDH-related parameters contributed substantially more to variance in fermentation flux than their CS-related counterparts over the chosen parameter ranges. However, total-order Sobol indices are unsigned and therefore do not indicate whether a parameter variation increases or decreases a given flux. We therefore examined the directional responses of the model using two experimentally motivated *in silico* redox perturbations. For each perturbation, we quantified the absolute TCA cycle and Pta–AckA fluxes, together with the fermentation ratio

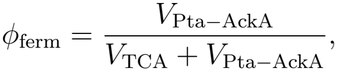

which represents the fraction of the modelled acetyl-CoA branch flux towards fermentation. Both perturbations were evaluated under four regulatory configurations: NADH inhibition of both PDH and CS, inhibition of PDH only, inhibition of CS only, and no inhibition. These labels refer to whether the corresponding inhibition factor was allowed to respond to changes in NADH. As described in Section 2.6, removing a response held its inhibition factor at the sample-specific baseline value, preserving baseline inhibition. The inhibitory interactions are shown by the red inhibitory arrows in Figure 2. We compared the four configurations using the one with both regulatory responses removed as the reference.

NADH oxidase overexpression was represented by decreasing the target redox ratio *α*. For each posterior sample, *α* was varied over a multiplicative range centred on its sample-specific posterior value *α*_*s*_, and the plotted trajectories were restricted to the common absolute range shared across the posterior ensemble.

We first examined how the PDH and CS regulatory responses altered the effect of varying the target redox ratio. Increasing *α* reduced TCA cycle flux under all four configurations (Figure 7A, left). This decline persisted when both inhibitory interactions were removed, showing that redox variation had a noticeable effect on pathway flux via the substrate availability alone. Retaining either the PDH or CS response amplified the decline, while retaining both produced the strongest reduction in TCA cycle flux. Thus, both regulatory responses further restricted TCA cycle throughput as *α* increased.

**Figure 7:**
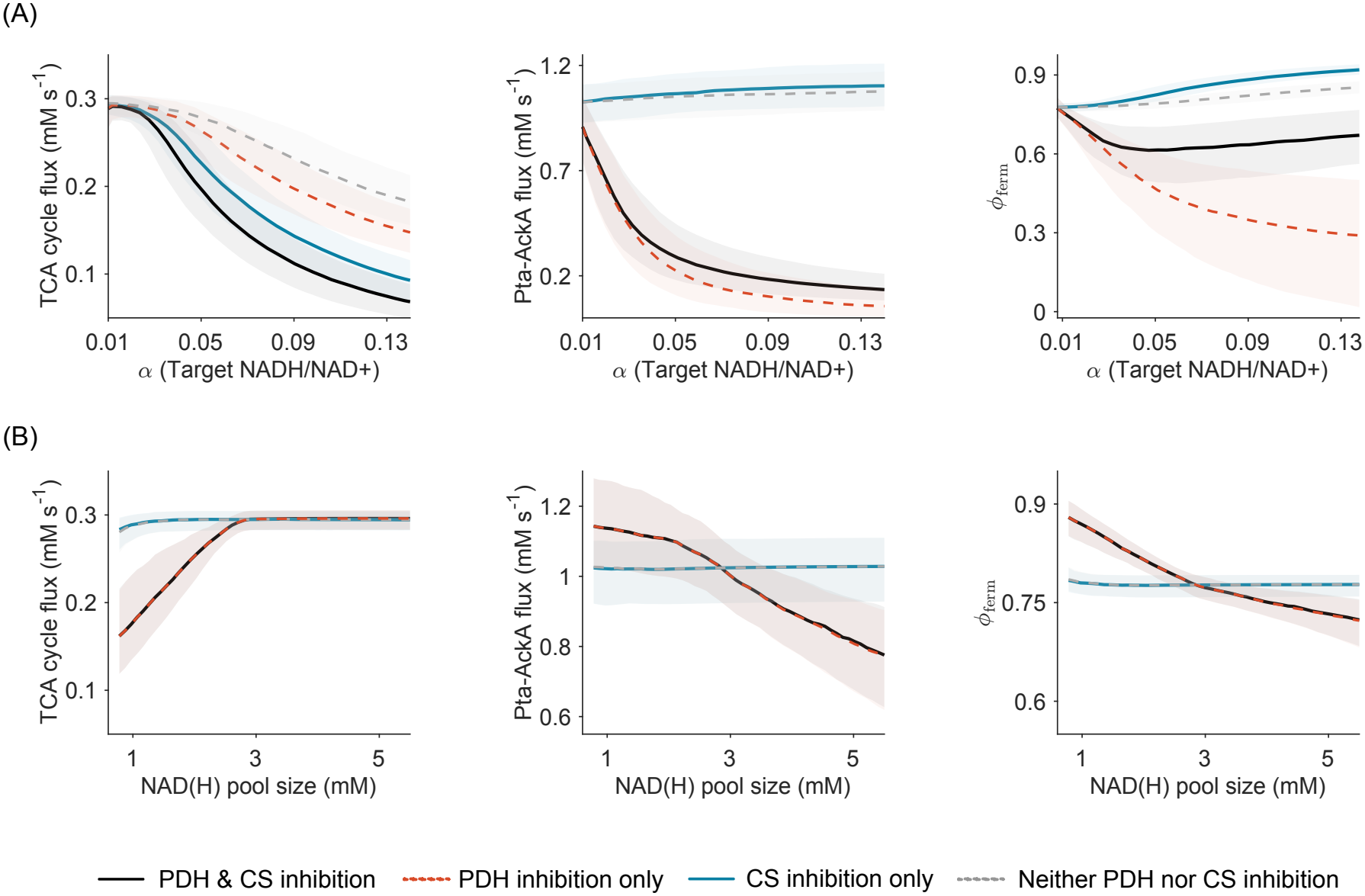
Steady-state flux and allocation responses to variation in (A) the target redox ratio *α* (NADH/NAD^+^) and (B) the total NAD(H) pool size *C*_NAD_. Columns show the TCA cycle flux, the absolute Pta–AckA flux, and the fermentation ratio *ϕ*_ferm_ = *V*_Pta*−*AckA_*/*(*V*_TCA_ + *V*_Pta*−*AckA_). Fluxes are reported in mM s^*−*1^ and the NAD(H) pool size in mM. Each perturbation was simulated under four regulatory configurations, with inhibition factors allowed to vary with NADH for both PDH and CS (solid black), PDH only (dashed orange), CS only (solid blue), or neither enzyme (dashed grey). Inhibition factors without NADH dependence were held at their sample-specific baseline values. Lines show the posterior medians of the steady-state values, and shaded regions show the corresponding central 50% posterior intervals.

The corresponding Pta–AckA responses depended primarily on whether PDH was regulated by NADH (Figure 7A, centre). When the PDH response was removed, Pta–AckA flux increased moderately with *α*, regardless of whether the CS response was retained. When the PDH response was retained, absolute Pta–AckA flux instead decreased strongly with increasing *α*, regardless of the CS response. This separation reflects the position of PDH upstream of the acetyl-CoA branch point: its NADH-dependent inhibition restricts acetyl-CoA supply and therefore reduces the absolute flux available to both the TCA cycle and the Pta–AckA pathway.

The fermentation ratio further distinguished the effects of the PDH and CS responses on relative carbon allocation (Figure 7A, right). With the PDH response removed, absolute Pta–AckA flux increased with *α* while TCA cycle flux declined, causing *ϕ*_ferm_ to increase. This increase was more pronounced when the CS response was retained, consistent with NADH-dependent CS inhibition preferentially restricting acetyl-CoA entry into the TCA cycle branch. With only the PDH response retained, Pta–AckA flux declined proportionally more than TCA cycle flux, and *ϕ*_ferm_ therefore decreased monotonically. When both responses were retained, *ϕ*_ferm_ initially decreased but partially recovered at higher *α*, as the decline in TCA cycle flux eventually became proportionally greater than that in Pta–AckA flux. Thus, the CS response favoured relative allocation towards fermentation, whereas the PDH response primarily restricted the absolute throughput supplied to both branches. Consequently, an increase in *ϕ*_ferm_ did not necessarily correspond to an increase in absolute Pta–AckA flux.

Together, these results distinguish regulation of pathway throughput from regulation of carbon allocation. The NADH-dependent PDH response primarily controlled acetyl-CoA supply and therefore absolute Pta–AckA flux, whereas the CS response influenced the relative partitioning of acetyl-CoA between the TCA cycle and fermentation. The effect of the CS response was most apparent in the fermentation ratio under the target-redox perturbation, even though CS-related parameters accounted for comparatively little variance in absolute fermentation flux in the broader Sobol analysis.

Vemuri et al. showed that overexpression of a water-forming NADH oxidase, which provides a non-respiratory route for NADH oxidation and lowers the NADH/NAD^+^ ratio, reduced acetate production and increased CO_2_ evolution at matched glucose uptake rates [10].

When compared with this experiment, all four regulatory configurations predicted that lowering *α* would increase TCA cycle flux (Figure 7A, left). Although TCA cycle flux is not a direct measure of CO_2_ evolution, this direction of change was qualitatively consistent with the reported increase in CO_2_ evolution. The direction of the Pta–AckA response, however, depended on the PDH response. With the PDH response removed, lowering *α* reduced Pta–AckA flux, consistent with the observed reduction in acetate production; retaining the CS response did not reverse this result. With the PDH response retained, lowering *α* instead increased Pta–AckA flux, opposite to the experimental observation. Thus, the NADH-dependent PDH response, rather than the CS response, prevented the fully regulated configuration from reproducing the acetate response to NADH oxidase overexpression.

### 3.4 PDH inhibition shapes flux allocation in response to total NAD(H) pool expansion

The elevated NAD(H) pool was represented by increasing the total pool size *C*_NAD_, while holding inferred parameters at their sample-specific posterior values and other fixed model quantities at their baseline values. We next used the same four regulatory configurations to examine how the PDH and CS regulatory responses altered the effect of expanding the total NAD(H) pool. With both regulatory responses removed, increasing the NAD(H) pool produced little change in TCA cycle flux, Pta–AckA flux, or *ϕ*_ferm_ (Figure 7B). When only the CS response was retained, these outputs remained approximately constant, indicating that the CS response contributed little to the pool expansion effect. In contrast, when the PDH response was retained, increasing the NAD(H) pool markedly increased TCA cycle flux, which approached a plateau at larger pool sizes, while reducing both absolute Pta–AckA flux and *ϕ*_ferm_. These responses were almost indistinguishable whether the CS response was retained or removed. Thus, the four configurations separated primarily according to whether the PDH response was retained, indicating that the predicted response to NAD(H)-pool expansion was mediated predominantly through the effective NADH response of PDH.

Han and Eiteman examined an *E. coli* strain with an elevated intracellular NAD(H) pool, generated by deleting the NAD(H)-degradation genes *nadR, nudC*, and *mazG*. In glucose-limited chemostat cultures, this strain produced less acetate at matched glucose uptake rates and exhibited a delayed onset of acetate overflow, which occurred only at a higher growth rate [50]. Despite the reduction in acetate production, the engineered and wild-type strains showed no significant differences in specific oxygen-consumption or CO_2_-evolution rates at matched glucose uptake rates [50].

Because the present model does not describe growth-rate-dependent overflow onset or respiration explicitly, we primarily compared the direction of the predicted acetate-flux response with the experimental observation. We also examined TCA cycle flux as a qualitative indicator of change in oxidative carbon metabolism, rather than as a direct prediction of oxygen consumption or CO_2_ evolution.

With the PDH response removed, the predicted TCA cycle flux remained approximately constant as the NAD(H) pool increased, qualitatively compatible with the lack of a significant change in the measured oxygen-consumption and CO_2_-evolution rates. However, Pta–AckA flux also remained approximately constant and therefore did not reproduce the observed reduction in acetate production. Retaining the PDH response produced the experimentally consistent decrease in Pta–AckA flux, but also a pronounced increase in TCA cycle flux. Although TCA cycle flux is not a direct measure of respiration or CO_2_ evolution, this predicted increase in oxidative pathway throughput was not matched by a detectable change in either experimental measure. Retaining or removing the CS response had little effect on these conclusions.

The two perturbations therefore imposed competing constraints on the effective NADH response of PDH. Retaining the full NADH-dependent PDH response generated the decrease in Pta–AckA flux following NAD(H)-pool expansion, but reversed the acetate-flux response to NADH oxidase overexpression and produced a pronounced pool-dependent increase in TCA cycle flux. Conversely, removing the PDH response recovered the experimentally consistent acetate-flux response to NADH oxidase overexpression and kept TCA cycle flux approximately constant during pool expansion, but eliminated the accompanying decrease in Pta–AckA flux.

### 3.5 A weak but non-zero PDH regulation reconciles experimentally observed responses to redox perturbations

These contrasting outcomes in Sections 3.3 and 3.4 suggested that the effective NADH response of PDH might need to be weaker than in the fully regulated model, but not absent altogether. We therefore tested whether a weak but non-zero response could preserve the decrease in Pta–AckA flux following NAD(H)-pool expansion, recover the acetate-flux response to NADH oxidase overexpression, and attenuate the degree to which PDH is regulated by NADH.

We introduced a phenomenological scaling parameter *β ∈* [0, 1] that continuously modulates the effective NADH response of PDH while preserving the calibrated baseline steady state. Here, *β* = 1 recovers the NADH dependence represented in the original model, whereas *β* = 0 holds this factor at its sample-specific baseline value, removing its response to changes in NADH concentration.

In the original model (Sections S1 and S1.2 in the Supplementary Materials), the PDH rate (Equation (30)) is given by

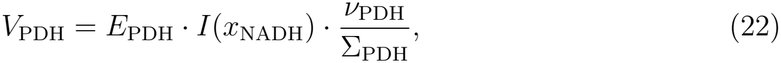

where the NADH inhibition factor is

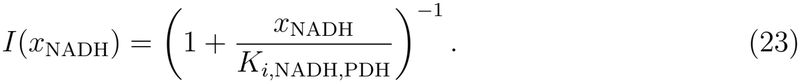

We rescale this inhibition factor through *β* as

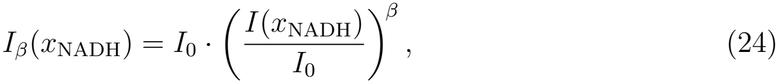

so that the PDH rate becomes

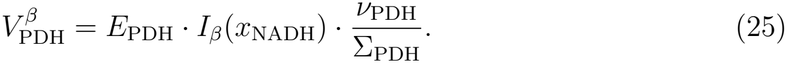

By construction, *I*_*β*_ = *I*_0_ at the baseline NADH concentration 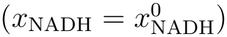 for every *β*, so that 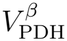 leaves the calibrated baseline steady state unchanged and only the response to deviations in *x*_NADH_ is rescaled.

The parameter *β* rescales the local log-sensitivity of the NADH inhibition term around baseline:

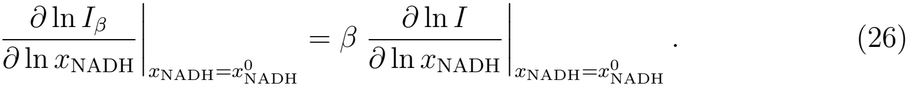

For each value of *β* and each posterior sample, we independently simulated the target-redox and NAD(H)-pool perturbations to steady state. We then fitted a linear function *V*_Pta*−*AckA_ = *c*_*α*_ + *m*_*α*_*f*_*α*_ by ordinary least squares, where *f*_*α*_ = *α/α*_0_ is the fold-change in *α* relative to that sample’s baseline value. For the pool perturbation, we analogously fitted *V*_Pta*−*AckA_ = *c*_pool_ + *m*_pool_*f*_pool_, where *f*_pool_ is the fold-change in the total NAD(H) pool relative to its baseline value.

The fitted intercepts were denoted *c*_*α*_ and *c*_pool_, and the corresponding response slopes were denoted *m*_*α*_ and *m*_pool_. At each *β*, the slopes were summarised by their posterior medians and interquartile ranges. The critical values 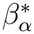 and 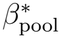 were defined by the zero-crossings of the corresponding posterior-median slopes.

At *β* = 1, the model predicted a negative response of fermentation flux to increasing *α*, contrary to the experimentally observed increase in acetate production, while retaining the negative response to increasing NAD(H)-pool size. As the effective PDH response was attenuated, *m*_*α*_ changed sign at 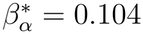 and became positive for smaller values of *β*. By contrast, reproducing the negative pool-size response required 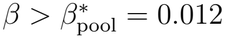. The two experimental trends were therefore simultaneously recovered only within the interval 0.012 *< β <* 0.104 (Figure 8B). Because *β* directly scales the local log-sensitivity of the NADH inhibition term, this interval corresponds to approximately 1.2% – 10.4% of the response represented in the original model. Thus, within the present model, the response directions reported for the two perturbations are simultaneously compatible with a weak but non-zero effective NADH response of PDH.

**Figure 8:**
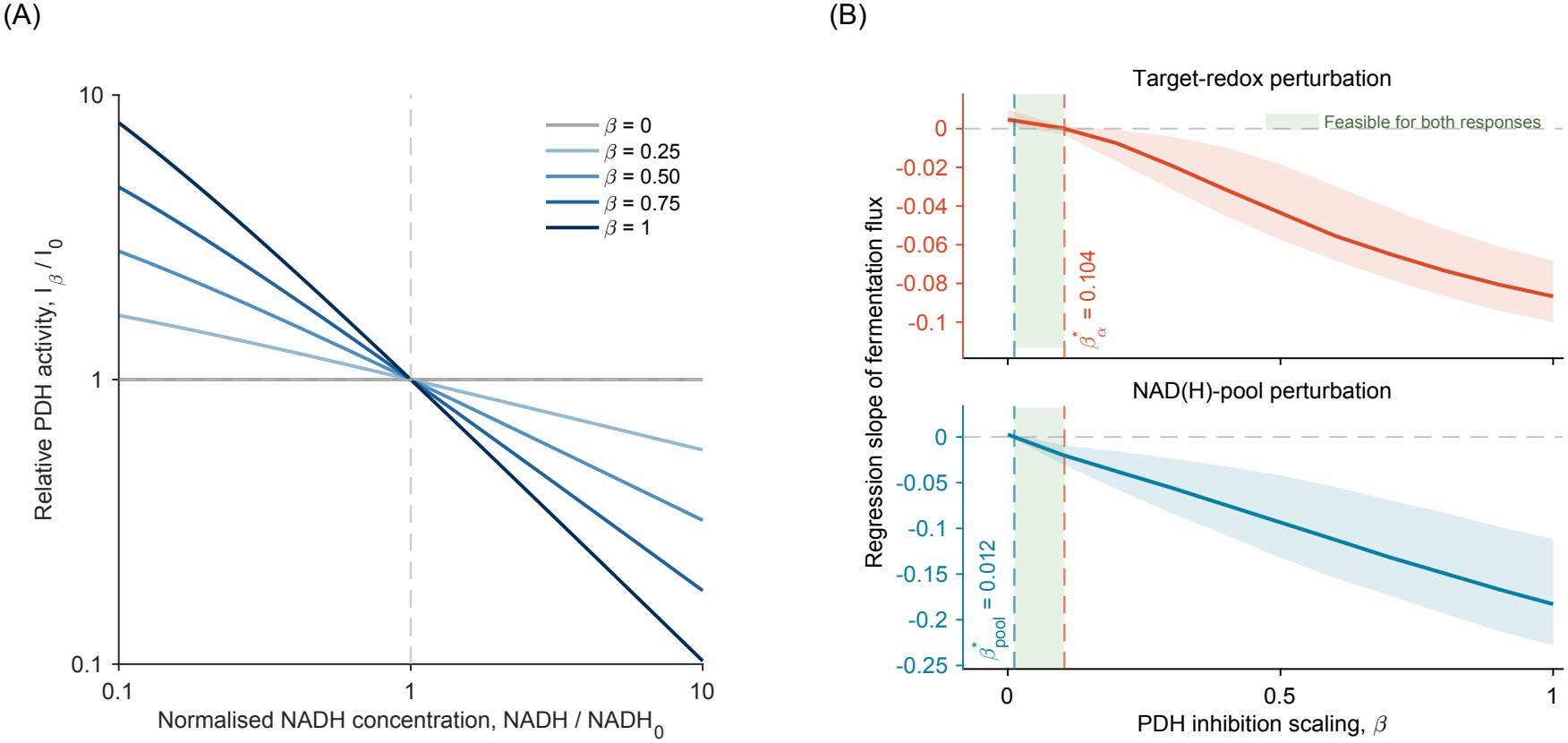
PDH NADH-response strengths compatible with the two redox perturbations. (A)Relative contribution of the NADH inhibition factor to PDH activity, *I*_*β*_*/I*_0_, as a function of normalised NADH concentration. All curves pass through the baseline point (1, 1), while decreasing *β* progressively attenuates the response to deviations from baseline. At *β* = 0, the inhibition factor is independent of NADH concentration. (B) Regression slopes of fermentation flux against the target-redox fold-change (orange, upper axis) and the NAD(H)-pool fold-change (blue, lower axis), plotted as functions of *β*. Solid lines show posterior medians and shaded bands show interquartile ranges. The horizontal dashed line indicates zero slope. Vertical dashed lines mark the zero-crossings 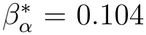 and 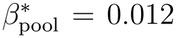. The experimentally observed responses require *m*_*α*_ *>* 0 and *m*_pool_ *<* 0, which are simultaneously satisfied for 0.012 *< β <* 0.104.

We examined the full perturbation responses at *β* = 0.05, a representative value within the compatible interval (Figure 9). With this attenuated but non-zero PDH response, increasing the target NADH/NAD^+^ ratio increased fermentation flux, consistent with the direction inferred from the NADH-oxidase experiment (Figure 9A). In contrast, increasing the total NAD(H) pool decreased fermentation flux, consistent with the response reported for the enlarged pool strain (Figure 9B). Thus, a single intermediate value of *β* reproduced the opposing fermentation responses to the two redox perturbations.

**Figure 9:**
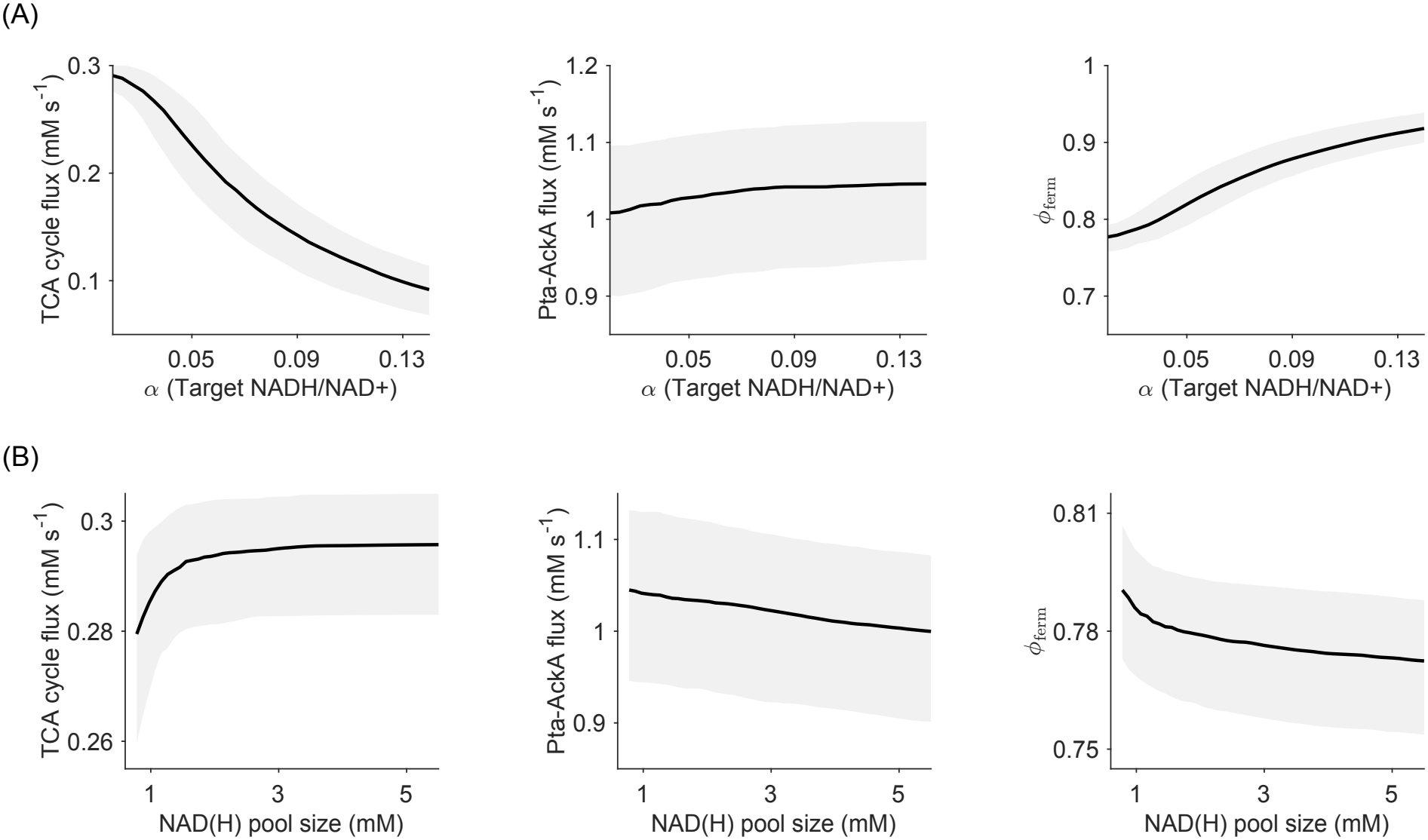
Steady-state fermentation responses to the two redox perturbations after attenuation of the effective NADH response of PDH. Predictions are shown for *β* = 0.05, which lies within the model-compatible interval identified from the two perturbation-response directions. Solid lines show posterior medians and shaded regions show the corresponding central 50% interval.

## 4 Discussion

This study investigated whether metabolite-dependent kinetics within the TCA cycle and fermentation pathways could translate a change in the cellular NAD(H) state into acetate-producing flux. Posterior predictive checks showed that the Bayesian-calibrated posterior ensemble was broadly consistent with the steady-state metabolite concentrations and fluxes reported by Park et al. [34] at the calibration condition (Figure 4). A strength of the present analysis is that these perturbations were evaluated across a Bayesian-calibrated posterior ensemble rather than using a single fitted parameter set, allowing uncertainty in the baseline kinetic model to propagate into the predicted responses.

In the global sensitivity analysis, the target-redox parameter *α* and the total NAD(H) pool influenced several model outputs. Among the enzyme-capacity parameters, Pta–AckA flux was most sensitive to *E*_PTA_, whereas *E*_AKGDH_ and *E*_SDH_ were among the strongest influences on TCA cycle flux. These enzyme rankings raise questions about how sensitivity within the reduced model relates to physiological flux control. In particular, the relatively strong effects of AKGDH and SDH on TCA cycle flux compared with those of CS, which has been described as a pace-making enzyme of the TCA cycle [51], warrant further investigation. Future work could assess their robustness across alternative calibrated parameter sets and perturbation ranges, compare them with local flux-control coefficients, and test the predicted influence of these enzymes through targeted changes in their abundances or activities under matched growth conditions.

Turning to the redox perturbation analysis, we used the calibrated posterior ensemble to examine perturbations inspired by reported changes in the NADH/NAD^+^ ratio [10] and in NAD(H) availability [50]. For the NAD(H)-pool perturbation, the full regulatory model predicted a decrease in Pta–AckA flux, qualitatively consistent with the lower acetate production reported by Han and Eiteman [50]. With the explicit NADH-inhibition response of PDH represented at its original strength, increasing the target-redox parameter reduced rather than increased Pta–AckA flux (Figure 7). However, attenuating this response allowed the model to reproduce the experimentally reported directions of both the target-redox and NAD(H)-pool perturbations [10, 50], provided that a weak residual response was retained (Figure 8).

Moreover, the redox total pool size and redox ratio perturbations provide complementary information because they probe different biochemical dimensions of the NAD(H) system. Altering the target-redox parameter changes the distribution of a fixed NAD(H) pool between its oxidised and reduced forms and therefore affects NAD^+^ substrate availability, NADH product effects, thermodynamic driving forces, and the encoded inhibitory interactions. Increasing the total NAD(H) pool instead changes the absolute availability of both cofactors and the capacity of the pool to buffer redox-dependent reactions; it is not equivalent to changing the NADH/NAD^+^ ratio. Consequently, the two perturbations need not produce the same flux response. In the model, the target-redox perturbation placed an upper bound on the explicit PDH response, because a stronger response caused the reduction in acetyl-CoA production to dominate (Figure 8). The pool-size perturbation placed a lower bound on the same response, because its experimentally consistent decrease in Pta–AckA flux was lost when the explicit PDH response became too weak (Figure 8). More broadly, this distinction indicates that the NADH/NAD^+^ ratio alone is insufficient to characterise the metabolic consequences of perturbing the NAD(H) system: redox state and redox pool capacity are related but non-equivalent control variables.

Within the scope of the model, we found that the direction of the fermentation response was not determined by the redox state alone, but by the balance between processes controlling acetyl-CoA supply and those controlling its consumption by the TCA and Pta–AckA branches. PDH therefore is hypothesised to control the total flux supplied to the modelled acetyl-CoA branch, whereas CS more directly affects how that supply is divided between TCA-associated oxidation and acetate production. Strong NADH inhibition of PDH can reduce the total acetyl-CoA production sufficiently that absolute Pta–AckA flux falls, even when the fraction of the remaining branch flux allocated towards Pta–AckA increases. In the full regulatory model, the TCA flux could fall proportionally more than the Pta–AckA flux, producing an increase in fermentation ratio, *ϕ*_ferm_, without an increase in absolute acetate-producing flux (Figure 7). Attenuating the explicit PDH response reduces this throughput penalty, allowing the allocation shift towards Pta–AckA to dominate the absolute fermentation response. This explains why the model can capture a relative shift towards fermentation while still incorrectly predicting the change in absolute Pta–AckA flux when PDH inhibition is too strong. These results therefore identify the relative strengths of competing regulatory effects at the acetyl-CoA node, rather than NADH accumulation alone, as a determinant of the acetate-flux response. Because upstream glycolytic supply and acetate exchange are fixed, the present model cannot test whether these local effects reproduce the glucose-uptake threshold for net acetate secretion observed experimentally [10]. Explicit regulation of the Pta–AckA branch and acetate reassimilation could also influence the onset of net secretion and warrant investigation in a broader model.

The range of *β* values obtained in Section 3.5 should not be interpreted as a direct estimate of intracellular PDH inhibition or of an effective intracellular *K*_*i*_. The parameter *β* rescales the responsiveness of the explicit NADH-inhibition factor while preserving its contribution at the calibrated baseline state. Moreover, even at *β* = 0, the reversible PDH rate law retains its dependence on NAD^+^ as a substrate and NADH as a product. The compatible interval therefore refers specifically to the log-sensitivity encoded by the explicit inhibition term, rather than to the total sensitivity of PDH flux to the NAD(H) state. Because the interval was determined from the directions of perturbation-response slopes rather than by quantitatively fitting the two experiments, it should be regarded as a model-conditional compatibility region, not as a confidence or credible interval for an intracellular biochemical parameter. Several biological effects could make the effective intracellular response weaker than that implied by transferring an isolated-enzyme inhibition term directly into the model. Accumulation of some other metabolites may also activate PDH and partially compensate for inhibition.

For example, fructose-1,6-bisphosphate and several other glycolytic intermediates stimulate the purified *E. coli* PDH complex [52]. Intracellular pH, ionic conditions, cofactor availability, macromolecular interactions, and additional metabolic effectors may also differ substantially from the conditions used to characterise purified enzymes. In addition, changes in PDH abundance or in competing pyruvate-consuming pathways could alter the system-level response without changing the inhibition of an individual PDH complex. However, these do not provide a direct quantitative mapping to *β*, which absorbs the net effect of multiple processes omitted from the reduced model.

We focused on local redox-dependent kinetics because NAD(H) directly participates in the reactions and regulatory interactions represented in the modelled pathways. These results position local redox-dependent kinetics as one layer of a broader, potentially complementary explanation of acetate overflow. Proteome-allocation constraints, respiratory or membrane-capacity limitations, coupling between ATP demand and NAD^+^ regeneration, and ArcA/ArcB-dependent transcriptional regulation may help determine when and why the intracellular redox state changes as glucose uptake and growth rate increase [3, 5, 6, 10]. The present model does not predict the origin of this redox imbalance.

Moreover, agreement with the Park et al. calibration data establishes consistency at one baseline condition rather than independent validation of the perturbation mechanism. The comparison with the NADH-oxidase experiment also requires caution. Expression of a heterologous NADH oxidase does not constitute a purely metabolic clamp of the NADH/NAD^+^ ratio. Vemuri et al. reported accompanying changes in the expression of TCA cycle and respiratory genes, including responses associated with ArcA regulation, as well as broader changes in cellular physiology [10]. The observed acetate response therefore cannot be interpreted as a direct measurement of allosteric regulation at fixed enzyme abundance. Similarly, enlargement of the NAD(H) pool in the strain studied by Han and Eiteman was produced through multiple gene deletions and examined after adaptation to steady growth [50], allowing regulatory changes beyond cofactor-pool expansion to contribute. Our model should therefore be understood as identifying the local kinetic conditions under which these intact-cell response directions are compatible with regulation within the TCA cycle and fermentation. It shows that the observed directions can arise without explicitly representing redox-responsive transcriptional regulation, but it does not demonstrate that transcriptional regulation was absent in either experiment.

Because enzyme abundances were held fixed, the perturbations represent steady-state metabolic adjustments under an expression-frozen parameterisation. The model does not explicitly simulate either the transient acute response or the slower adaptation of enzyme expression. In growing cells, however, acetate overflow is accompanied by growth-rate-dependent reallocation of glycolytic, TCA cycle, Pta–AckA, anaplerotic, and respiratory enzymes. Proteomic measurements indicating that PDH abundance remains stable or may increase across relevant growth-rate ranges [53] are particularly relevant: increased enzyme abundance or substrate supply could partly offset inhibition of activity at the level of each enzyme complex. The long-term in vivo response is therefore likely to reflect the combined effects of acute metabolic regulation and slower adaptation of enzyme expression, rather than either acting alone.

The restricted model boundary used in this paper limits the interpretation of the perturbation responses. Upstream glycolytic supply and downstream acetate exchange are represented through fixed boundary conditions, allowing the analysis to isolate regulation within the TCA cycle-fermentation module but excluding potential compensation through changes in glycolytic substrate supply. The effective PDH response identified here is therefore conditional on this reduced representation and may partly reflect influences that would emerge explicitly in a broader central carbon metabolism model. Similarly, the predicted Pta–AckA flux represents acetate production rather than a complete prediction of net extracellular acetate accumulation, which may also depend on acetate exchange and reassimilation. Comparisons with the experimental acetate responses should therefore be interpreted primarily at the level of response direction rather than quantitative secretion rate. Future extensions that make upstream carbon supply and extracellular acetate exchange dynamic would allow these contributions to be tested explicitly. Acute redox perturbations combined with altered PDH or citrate-synthase NADH sensitivity would be particularly informative for separating immediate enzyme-level regulation from subsequent transcriptional adaptation.

## 5 Conclusions

In conclusion, the model does not establish local NADH regulation as the sole cause of acetate overflow. Rather, it shows that the modelled pathways may have an enzyme-level regulatory capacity to translate redox perturbations into increased acetate-producing fluxes. However, this capacity is realised only when the explicit NADH response of PDH is sufficiently weak so that it does not suppress total acetyl-CoA production, while remaining strong enough to account for the NAD(H)-pool response. The association between redox imbalance and acetate overflow therefore depends not only on the redox state itself, but on how that state is transmitted through competing controls over pathway throughput and branch allocation. This leads to two nested questions for future work: what generates redox imbalance as carbon uptake and growth increase, and under what conditions, and through which metabolic and regulatory mechanisms, is that imbalance translated into acetate overflow?

## Supplementary Materials

### S1 Model structure and assumptions

#### S1.1 Notation

Let *x*_*q*_ denote the concentration of metabolite *q* (mM), and let *E*_*p*_ denote the concentration of enzyme *p* (mM). For convenience, metabolite concentrations are normalised by their reaction-specific Michaelis constants:

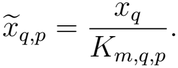

In this paper, *K*_*m,q,p*_ denotes the Michaelis constant of metabolite *q* for the reaction catalysed by enzyme *p*, and *K*_*i,q,p*_ denotes the inhibition constant of inhibitor *q* acting on enzyme *p*, both with units of mM. The parameters 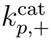 and 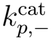 denote the forward and reverse catalytic turnover numbers (s^*−*1^), respectively.

Parameter values were collected from the BRENDA database and primary literature sources, the original and balanced parameter values were provided in Supplementary Table S1.

#### S1.2 Reaction rate laws

##### Pyruvate kinase

Pyruvate kinase (EC 2.7.1.40) catalyses the reaction

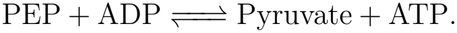

The reaction rate is represented by

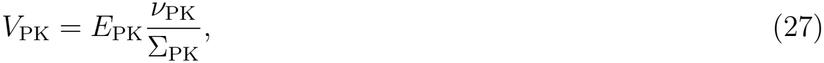

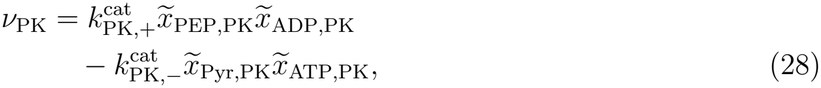

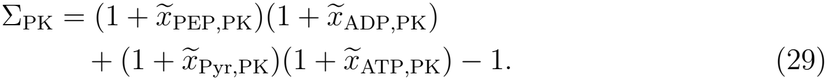

##### Pyruvate dehydrogenase

Pyruvate dehydrogenase (EC 1.2.4.1) catalyses the reaction

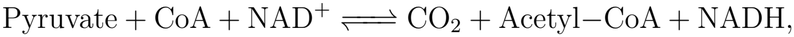

and is inhibited by NADH [11]. The reaction rate is represented by

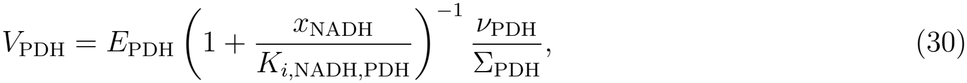

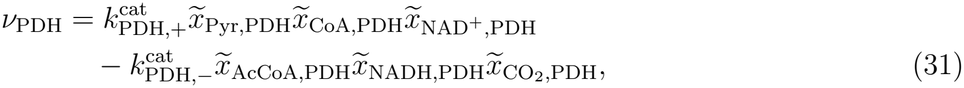

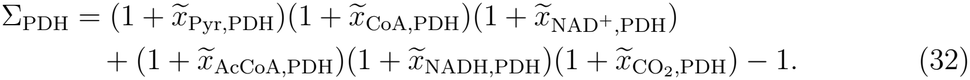

##### Citrate synthase

Citrate synthase (EC 2.3.3.1) catalyses the reaction

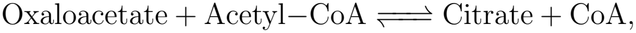

which is inhibited by NADH [12, 54]. The reaction rate is represented by

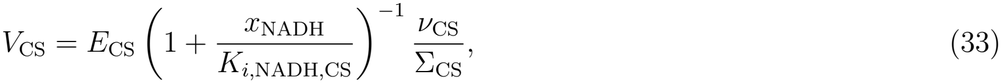

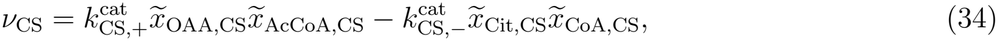

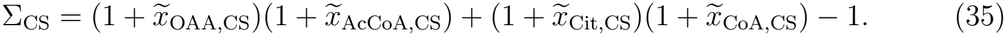

##### Aconitase

Aconitase (EC 4.2.1.3) catalyses the reaction

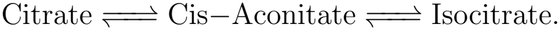

The rates of this two step reaction are expressed by

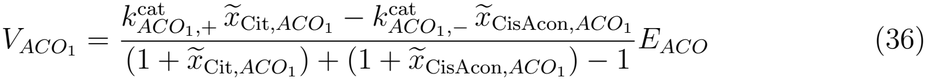

and

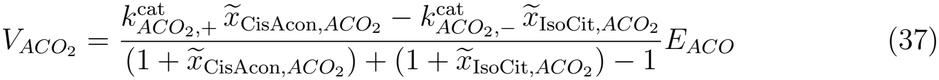

respectively.

##### Isocitrate dehydrogenase

Isocitrate dehydrogenase (EC 1.1.1.41) catalyses the reaction

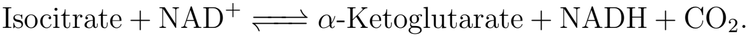

The reaction rate is represented by

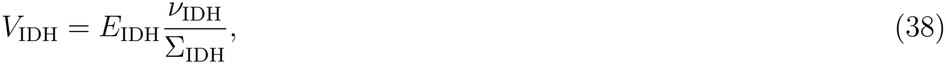

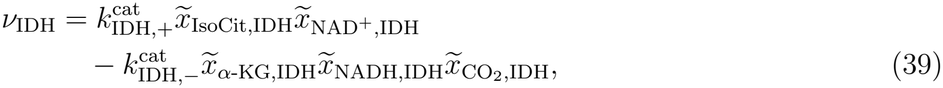

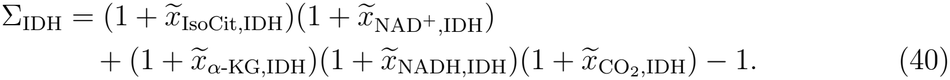

##### *α*-Ketoglutarate dehydrogenase

The *α*-ketoglutarate dehydrogenase catalyses the reaction of

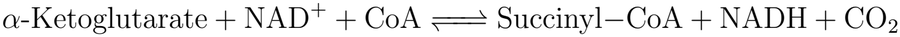

at the rate

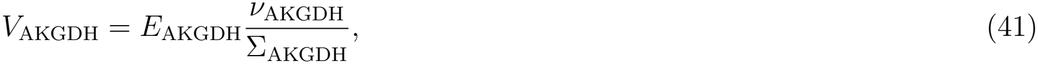

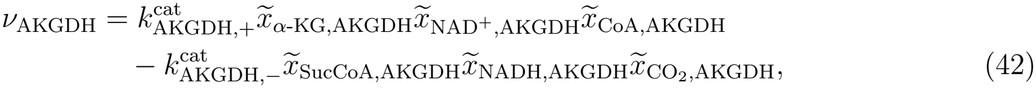

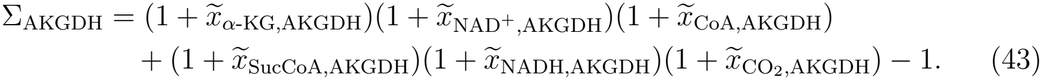

##### Succinyl-CoA synthetase

Succinyl-CoA synthetase (EC 6.2.1.5) catalyses the reaction

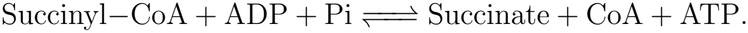

The reaction rate is represented by

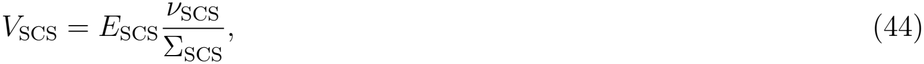

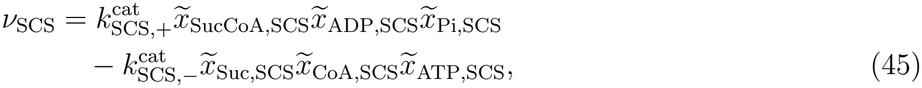

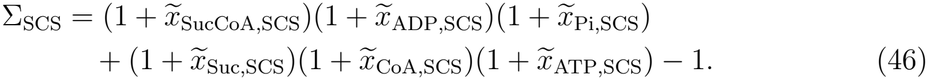

##### Succinate dehydrogenase

Succinate dehydrogenase (EC 1.3.5.1) catalyses the reaction

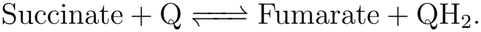

The reaction rate is represented by

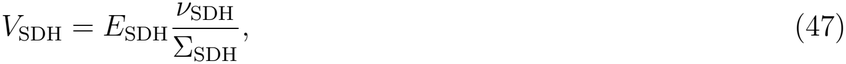

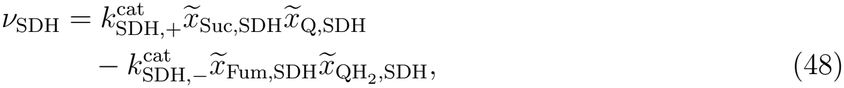

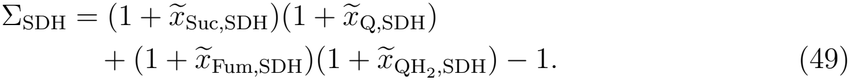

##### Fumarase

Fumarase (EC 4.2.1.2) catalyses the reaction

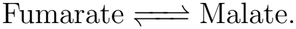

The reaction rate is represented by

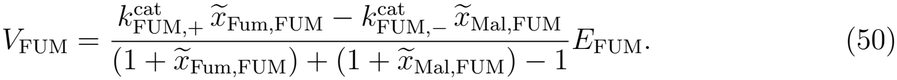

##### Malate dehydrogenase

Malate dehydrogenase (EC 1.1.1.37) catalyses the reaction

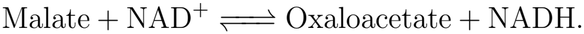

The reaction rate is represented by

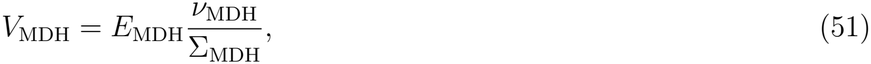

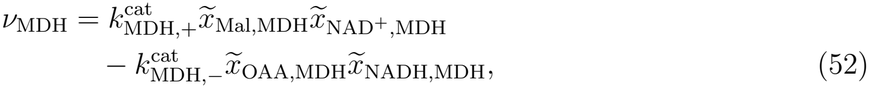

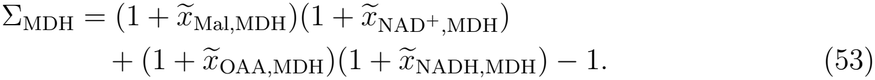

##### Phosphate acetyltransferase

Phosphate acetyltransferase (EC 2.3.1.8) catalyses the reaction

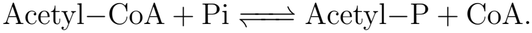

The reaction rate is represented by

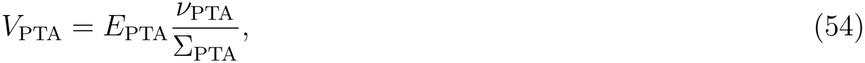

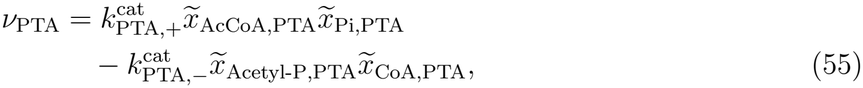

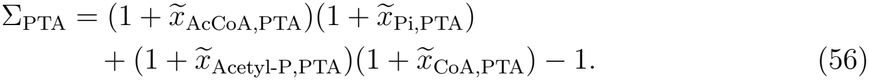

##### Acetate Kinase

Acetate kinase (EC 2.7.2.1) catalyses the reaction

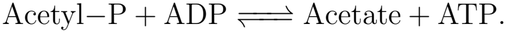

The reaction rate is represented by

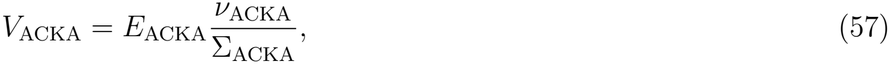

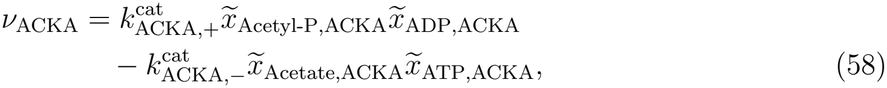

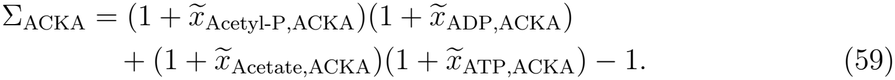

#### S1.3 Assumptions

##### Boundary variables

To focus on the TCA cycle and acetate fermentation branch, selected metabolites (PEP, ATP, ADP, CO_2_, Q, QH_2_, Pi, and acetate) were treated as boundary variables with fixed concentrations. Their values were fixed to represent external physiological conditions and coupling to the surrounding metabolic network, including source and sink reactions.

##### Phenomenological redox conversion

The full set of NAD^+^-regenerating and NADH-consuming reactions is outside the model network and not explicitly represented. To capture the dynamic interconversion of the NAD(H) pool, we introduced two lumped phenomenological processes representing the reduction of NAD^+^ and the oxidation of NADH, respectively:

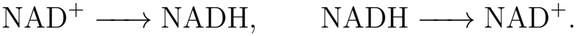

The corresponding directional rates are

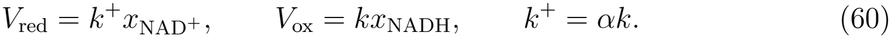

Their net contribution to the NADH balance is therefore

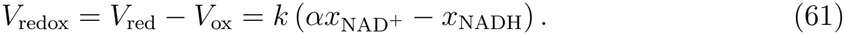

This phenomenological boundary representation is not thermodynamically consistent because it aggregates externally driven NADH-producing and NADH-consuming processes, and represents these as irreversible reactions.

We treat *k* and *α* as free parameters. The parameter *k* is the effective first-order rate constant for NADH oxidation and hence NAD^+^ regeneration, with units of s^*−*1^, whereas *α* defines the target NADH/NAD^+^ ratio towards which the phenomenological redox-conversion module drives the system. In the limit *k → ∞*, the interconversion acts as a rapid redox clamp, recovering the fixed boundary-ratio assumption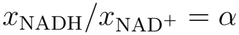. For finite *k*, the steady-state ratio can deviate from *α* depending on the balance of NADH production and consumption by the other reactions in the model.

##### Inhibition terms

In the present model, we only incorporate regulatory mechanisms with putative effects on the redox state, as these are the primary mechanisms of interest in our study. Other regulatory influences are not modelled individually; instead, their net effects are assumed to be absorbed into the effective kinetic parameters. Accordingly, regulatory terms are only introduced for selected gatekeeping enzymes of pyruvate dehydrogenase and citrate synthase [11, 12, 54], rather than for every weakly regulated reaction; this is to focus the model on the control points that couple carbon partitioning to redox state.

##### Conserved pools

Conservation laws arise from the stoichiometric structure of the network and correspond to linear combinations of metabolite concentrations that remain constant over time [35]. In general, any nonzero constant vector ***ℓ*** satisfying

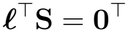

defines a conservation law. It follows that

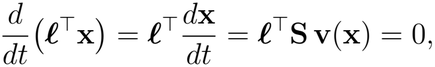

so that ***ℓ***^*⊤*^**x**(*t*) remains invariant over time [35].

The stoichiometric matrix **S** is constructed directly from the reaction network (Equations (1) to (15) in the main text).

|  | PK | PDH | CS | ACO<sub>1</sub> | ACO<sub>2</sub> | IDH | AKGDH | SCS | SDH | FUM | MDH | PTA | ACKA | Redox |
|---|---|---|---|---|---|---|---|---|---|---|---|---|---|---|
| Pyr | 1 | −1 | 0 | 0 | 0 | 0 | 0 | 0 | 0 | 0 | 0 | 0 | 0 | 0 |
| OAA | 0 | 0 | −1 | 0 | 0 | 0 | 0 | 0 | 0 | 0 | 1 | 0 | 0 | 0 |
| AcCoA | 0 | 1 | −1 | 0 | 0 | 0 | 0 | 0 | 0 | 0 | 0 | −1 | 0 | 0 |
| Cit | 0 | 0 | 1 | −1 | 0 | 0 | 0 | 0 | 0 | 0 | 0 | 0 | 0 | 0 |
| CisAcon | 0 | 0 | 0 | 1 | −1 | 0 | 0 | 0 | 0 | 0 | 0 | 0 | 0 | 0 |
| IsoCit | 0 | 0 | 0 | 0 | 1 | −1 | 0 | 0 | 0 | 0 | 0 | 0 | 0 | 0 |
| α-KG | 0 | 0 | 0 | 0 | 0 | 1 | −1 | 0 | 0 | 0 | 0 | 0 | 0 | 0 |
| SucCoA | 0 | 0 | 0 | 0 | 0 | 0 | 1 | −1 | 0 | 0 | 0 | 0 | 0 | 0 |
| Succ | 0 | 0 | 0 | 0 | 0 | 0 | 0 | 1 | −1 | 0 | 0 | 0 | 0 | 0 |
| Fum | 0 | 0 | 0 | 0 | 0 | 0 | 0 | 0 | 1 | −1 | 0 | 0 | 0 | 0 |
| Mal | 0 | 0 | 0 | 0 | 0 | 0 | 0 | 0 | 0 | 1 | −1 | 0 | 0 | 0 |
| CoA | 0 | −1 | 1 | 0 | 0 | 0 | −1 | 1 | 0 | 0 | 0 | 1 | 0 | 0 |
| Acetyl − P | 0 | 0 | 0 | 0 | 0 | 0 | 0 | 0 | 0 | 0 | 0 | 1 | −1 | 0 |
| NAD<sup>+</sup> | 0 | −1 | 0 | 0 | 0 | −1 | −1 | 0 | 0 | 0 | −1 | 0 | 0 | −1 |
| NADH | 0 | 1 | 0 | 0 | 0 | 1 | 1 | 0 | 0 | 0 | 1 | 0 | 0 | 1 |

with rows corresponding to metabolites and columns corresponding to reaction fluxes. In this model, the left null space of ***S*** is three-dimensional, and implies the following conserved quantities:

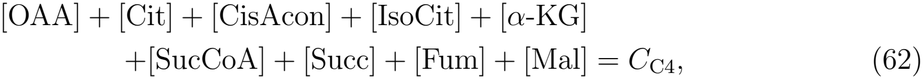

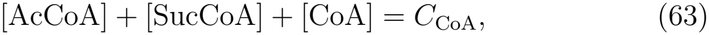

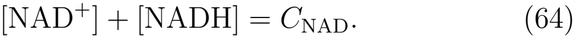

The values of these conserved quantities were set equal to the sums of the corresponding experimentally reported metabolite concentrations from Park et al. [34], as described in Section S2.1 in Supplementary Material and listed in Supplementary Table S1.

### S2 Parameterisation and data sources

#### S2.1 Data sources and parameter balancing method

Experimental data used for model calibration were obtained from Park et al. [34], who reported intracellular metabolite concentrations measured by liquid chromatography mass spectrometry (LC–MS) and steady-state fluxes estimated using steady-state ^13^C metabolic flux analysis (MFA) in *Escherichia coli* grown aerobically in glucose minimal medium. Only metabolite concentrations and fluxes corresponding to reactions or state variables represented in the reduced model were considered for calibration.

Our model does not represent anaplerotic inputs or carbon withdrawal towards biosynthetic pathways. Its steady-state mass balances therefore require equal net rates through the sequential TCA-cycle reactions. In contrast, the reaction-specific fluxes reported by Park et al. were not identical around the cycle because these branch fluxes were included in the full MFA network. We therefore imposed a common steady-state TCA-cycle throughput and used the minimum reported internal TCA-cycle flux, approximately 18 mM min^*−*1^, as an operational estimate of its value. This value represents the flux completing the cycle in the reduced model, without the contributions associated with the omitted anaplerotic and biosynthetic branches. For the fermentation branch, the aggregate EX AC flux reported by Park et al., which includes acetate production and acetyl-CoA demand for fatty-acid synthesis, was used as a proxy for Pta–AckA flux. Fatty-acid synthesis was not represented explicitly in the reduced model. Because Pta and AckA are represented as sequential reactions and no alternative acetyl-phosphate or acetate-producing routes are included, the steady-state balance requires *v*_Pta_ = *v*_AckA_. The reported EX_AC_ flux was therefore mapped to this common Pta–AckA pathway flux. The original reaction-specific measurements and their mapping to the reduced model are summarised in Supplementary Table S2.

For numerical simulations, initial values were used as starting points for integration toward steady state. Experimentally reported concentrations were used where available, while unmeasured state variables were initialised based on model assumptions. Boundary variables were fixed to experimentally reported values where available; otherwise, they were treated as unknown quantities and inferred during model calibration.

Kinetic constants are usually measured experimentally *in vitro*, but available values are often uncertain, incomplete, and inconsistent across studies [55]. Parameter balancing provides a systematic framework for integrating such kinetic data into complete and thermodynamically consistent parameter sets, while accounting for predefined parameter ranges and enforcing Wegscheider conditions and Haldane relationships [55]. Haldane relationships link forward and reverse catalytic constants and Michaelis constants to reaction equilibrium constants, while Wegscheider conditions ensure consistency of equilibrium constants across stoichiometrically closed reaction cycles. In this study, kinetic data, including Michaelis constants, catalytic rate constants, and inhibition constants, were collected from BRENDA (Supplementary Table S1) [32]. Equilibrium constants were obtained from eQuilibrator [33] using the default settings: pH 7.5, pMg 3.0, an ionic strength of 0.25 M, and a temperature of 298.15 K. These data were used as input to the online parameter balancing tool developed by Lubitz et al. [30], which combines available measurements with prior distributions and thermodynamic constraints to estimate a complete, thermodynamically consistent parameter set (Supplementary Table S1).

#### S2.2 Structural identifiability analysis

Structural identifiability was assessed using GenSSI before practical identifiability analysis and parameter estimation. The initial candidate parameter set comprised the 12 effective enzyme activity parameters, the redox parameters *k* and *α*, and the unknown boundary concentrations *x*_Acetate_, *x*_*Q*_, and 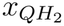. The remaining kinetic constants, including Michaelis constants and catalytic rate constants, were fixed at their parameter-balanced values. Inhibition constants and known boundary metabolite concentrations were fixed at their literature values. The output vector used for the structural identifiability analysis was

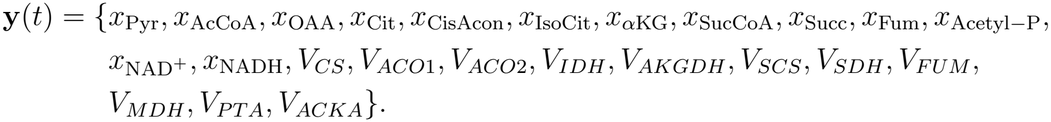

Initial conditions were treated as known. GenSSI assumes that these outputs are observed continuously and without measurement noise. Consequently, the analysis represents a best-case assessment of parameter identifiability under ideal dynamic observations, rather than a direct assessment of identifiability from the finite steady-state data used in the subsequent calibration.

Analysis of the full candidate parameter set showed that the complete parameter vector was not structurally identifiable. Most parameters were structurally globally identifiable, *E*_SDH_, *E*_ACKA_, and *x*_Acetate_ were structurally locally identifiable, and *x*_*Q*_ and 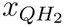 were structurally non-identifiable. The non-identifiable boundary concentrations *x*_*Q*_ and 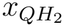 were therefore fixed at their assumed physiological values, and the analysis was repeated. After fixing *x*_*Q*_ and 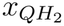 , no parameters remained structurally non-identifiable. The parameters

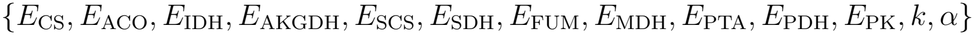

were structurally globally identifiable, whereas *x*_Acetate_ and *E*_ACKA_ remained structurally locally identifiable. Thus, although each of these two parameters was locally distinguishable, the model admitted a finite set of alternative solutions for the parameter pair.

Fixing either *x*_Acetate_ or *E*_ACKA_ resolved the remaining ambiguity and rendered all other parameters structurally globally identifiable. Because acetate is represented as a boundary condition, whereas *E*_ACKA_ represents the effective catalytic capacity of the acetate-fermentation branch, *x*_Acetate_ was fixed at the baseline boundary value used in the model simulations. The analysis was then repeated with *E*_ACKA_ retained as an unknown parameter, after which all remaining parameters were structurally globally identifiable.

The final parameter vector retained for practical identifiability analysis was therefore

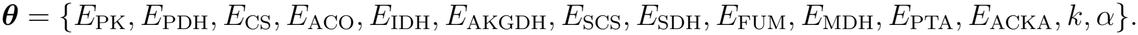

The successive reductions of the candidate parameter set are summarised in Table 1.

#### S2.3 Synthetic-data recovery study

Synthetic data were generated to evaluate whether the calibration workflow could recover known parameter values under the assumed model structure. A reference parameter vector, denoted by ***θ***^*\**^ (listed in Table 2), was used to simulate the kinetic model to steady state under the same parameter conditions as the experimental calibration. The resulting steady-state metabolite concentrations and fluxes were treated as the noise-free model outputs.

**Table 2:** Ground-truth values of the 14 model parameters in the reference parameter vector ***θ***^*\**^ used to generate the synthetic dataset for the parameter-recovery study. All values are reported on the original scale.

| Parameter | Ground-truth value | Unit |
|---|---|---|
| *E*<sub>PK</sub> | 0.1 | mM |
| *E*<sub>PDH</sub> | 0.1 | mM |
| *E*<sub>CS</sub> | 0.1 | mM |
| *E*<sub>ACO</sub> | 0.1 | mM |
| *E*<sub>IDH</sub> | 0.1 | mM |
| *E*<sub>AKGDH</sub> | 0.1 | mM |
| *E*<sub>SCS</sub> | 0.1 | mM |
| *E*<sub>SDH</sub> | 0.1 | mM |
| *E*<sub>FUM</sub> | 0.1 | mM |
| *E*<sub>MDH</sub> | 0.1 | mM |
| *E*<sub>PTA</sub> | 0.1 | mM |
| *E*<sub>ACKA</sub> | 0.1 | mM |
| *k* | 100 | s<sup>−1</sup> |
| α | 0.0195 | Dimensionless |

The *E* parameters represent effective enzyme abundances. The target redox parameter was set to α<sup>\*</sup> = 0.0545/2.8; the decimal value shown is rounded.

Synthetic observations were then generated by adding observation noise according to the same error model used in the calibration analysis. Metabolite concentrations were perturbed on the log_10_ scale, consistent with a multiplicative error model, while flux observations were generated using the same likelihood form as the experimental flux data (Equation (20), (21)). The resulting synthetic dataset therefore retained the observation structure of the experimental dataset, while providing known ground-truth parameter values. This “simulation-study” is an important process to confirm that the inference process is working: we generate synthetic data using some ***θ***^*\**^, then ‘forget’ ***θ***^*\**^ and attempt to reinfer it from the synthetic data [37].

#### S2.3.1 Practical identifiability analysis

Practical identifiability was assessed using profile-likelihood analysis. For each parameter *θ*_*k*_, the parameter was fixed at successive values within a predefined range, while the remaining parameters *θ*_*−k*_ were re-optimised by minimising the negative log-likelihood. The resulting profile negative log-likelihood was converted to a normalised profile log-likelihood difference according to

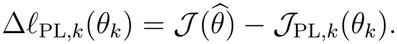

Thus, the normalised profile attained a maximum of zero at the maximum-likelihood estimate and decreased as *θ*_*k*_ was constrained away from its optimum. The approximate 95% likelihood-ratio threshold was

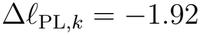

for a one-dimensional parameter profile [40]. The resulting profiles were used to assess whether the known parameter values could be recovered from the synthetic observations.

A parameter was classified as two-sided practically recoverable when its profile crossed the likelihood-ratio threshold on both sides of the maximum. Profiles crossing the threshold on only one side were classified as weakly practically identifiable, whereas profiles that failed to cross the threshold in either direction were classified as practically non-identifiable under the synthetic observation scheme [39, 56].

**Figure S1:**
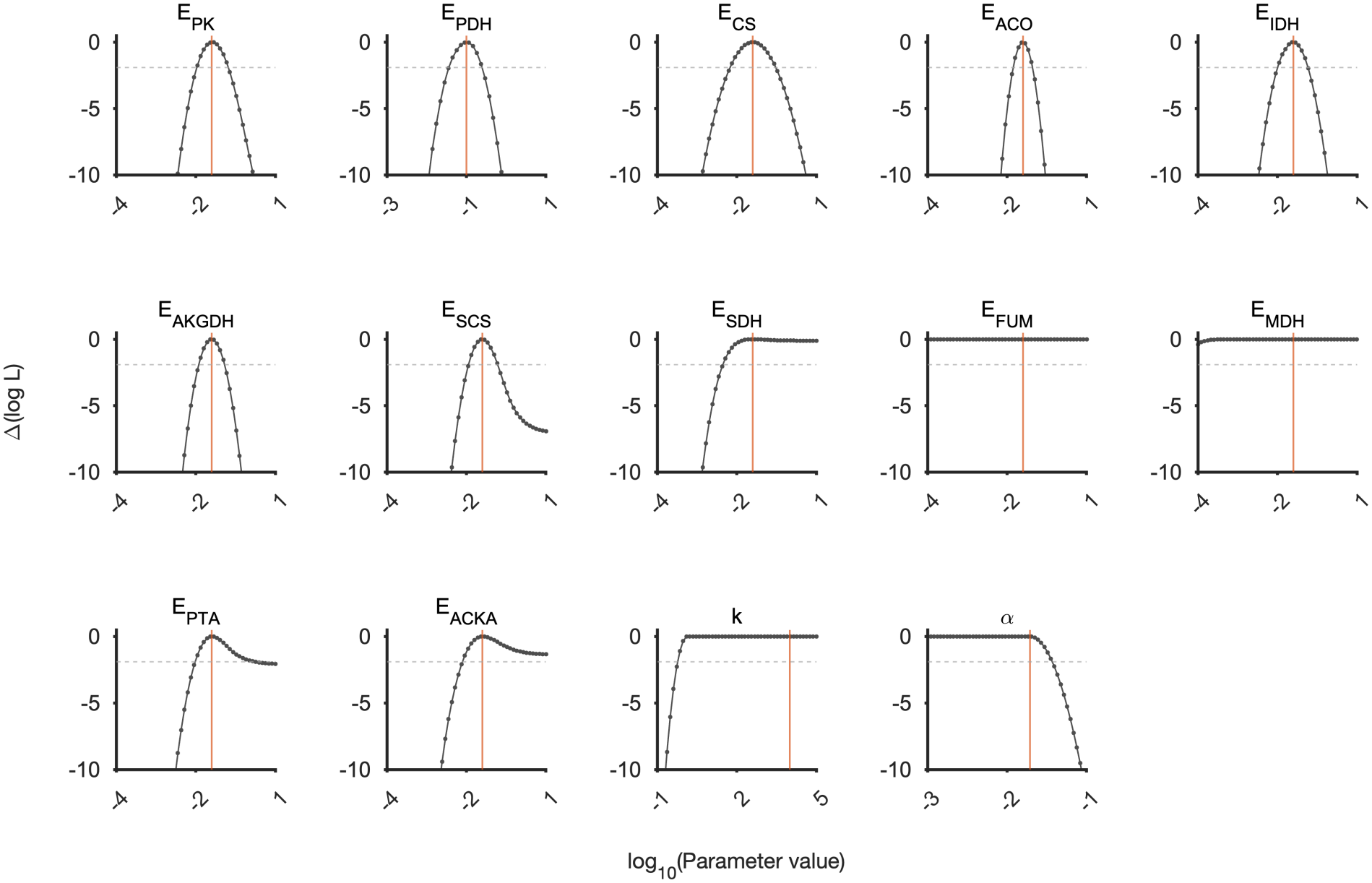
Normalised profile log-likelihoods for the parameters retained after structural identifiability analysis, calculated using synthetic steady-state observations. Each profile is shown relative to its maximum, such that Δℓ_PL_ = 0 at the maximum-likelihood estimate. The horizontal dashed line denotes the approximate 95% likelihood-ratio threshold, Δℓ_PL_ = *−*1.92. Vertical orange lines indicate the ground-truth parameter values used to generate the synthetic data.

The profiles for *E*_PK_, *E*_PDH_, *E*_CS_, *E*_ACO_, *E*_IDH_, *E*_AKGDH_, *E*_SCS_, and *E*_PTA_ crossed the likelihood-ratio threshold on both sides of their maxima, indicating two-sided recoverability under the synthetic observation scheme (Figure S1). Their profile maxima were also located close to the corresponding ground-truth values.

In contrast, *E*_SDH_, *E*_ACKA_, *k* and *α* were only one-sided recoverable and therefore weakly practically identifiable. The profiles for *E*_FUM_ and *E*_MDH_ remained approximately flat and did not cross the likelihood-ratio threshold, indicating practical non-identifiability from the synthetic steady-state observations (Figure S1). These results illustrate that parameters that are structurally globally identifiable under ideal dynamic observations may remain weakly constrained when only finite, noisy steady-state observations are available. The synthetic profile-likelihood analysis was used as a controlled assessment of parameter recovery under the assumed model and observation structure (Figure S1). It was not used to determine the final prior treatment for the experimental-data calibration; that treatment was based on the profile-likelihood analysis of the experimental observations described in Supplementary Section S2.4.1. Parameters with two-sided profiles were assigned broad priors, whereas parameters with one-sided or flat profiles were assigned independently informed priors based on available biochemical, proteomic, or physiological information. All parameters in the final structurally identifiable parameter set were retained so that uncertainty in weakly constrained but potentially influential parameters could be propagated to the model predictions.

#### S2.3.2 Bayesian estimation and posterior recovery

The synthetic dataset was then fitted using the Bayesian calibration workflow used in the main analysis. Posterior recovery was evaluated by comparing the inferred posterior distributions with the corresponding priors and the known ground-truth parameter values.

For most inferred parameters, the posterior distributions were concentrated relative to the priors and were centred near the ground-truth values (Fig. S2). Parameters with broader posterior intervals (*E*_*FUM*_ , *E*_*MDH*_ , *E*_*SDH*_ and *k*) were less strongly constrained by the available steady-state observations, consistent with weaker practical identifiability under the given observation scheme (Fig. S2).

**Figure S2:**
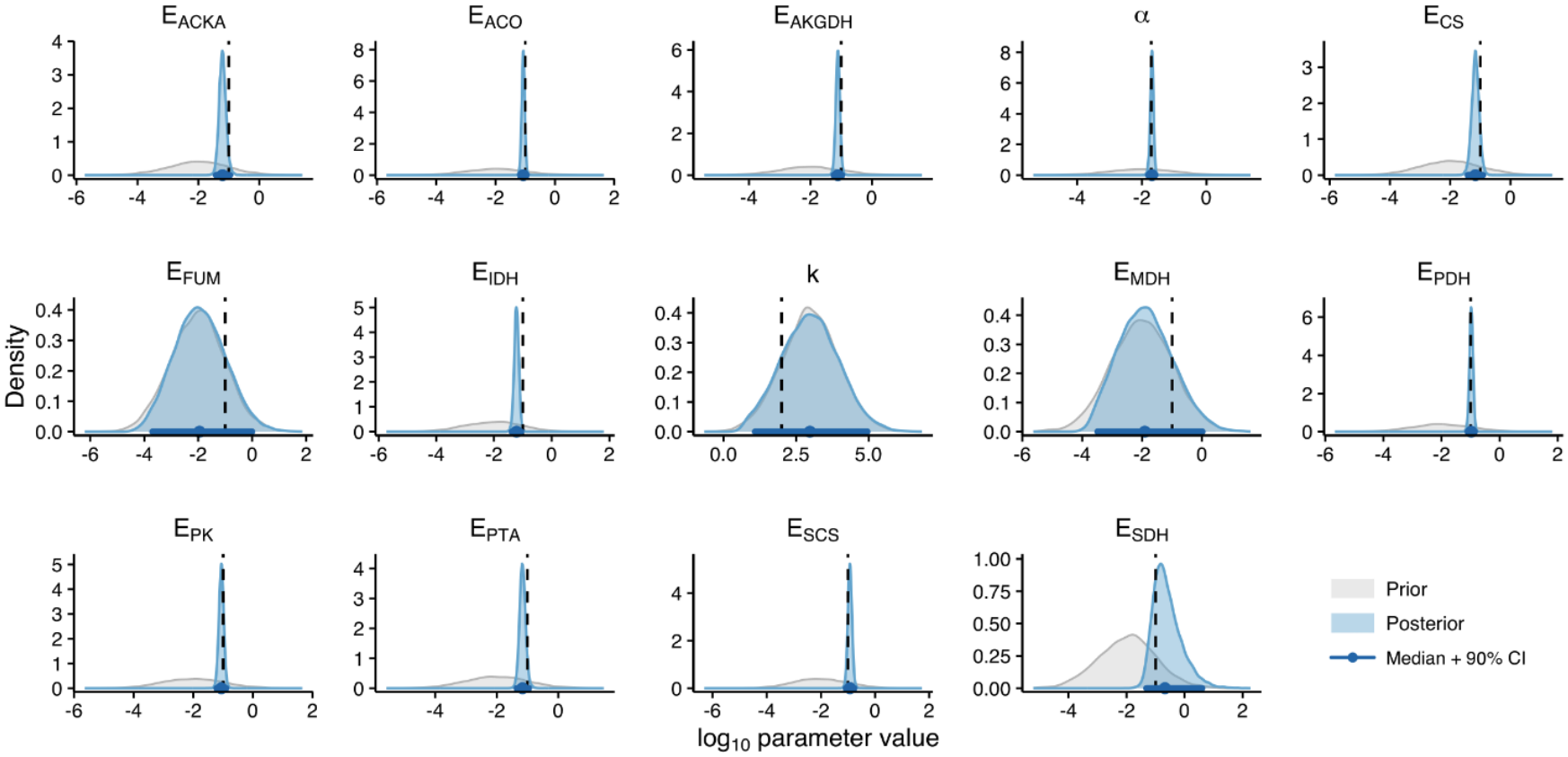
Prior–posterior comparison for inferred parameters in the synthetic-data recovery test. Grey densities show the prior distributions, blue densities show the posterior distributions, and black vertical dash lines indicate the ground-truth parameter values used to generate the synthetic data. Blue points and horizontal intervals indicate posterior medians and 90% credible intervals.

Posterior predictive checks showed that the fitted model reproduced the synthetic steady-state metabolite concentrations and fluxes, with the synthetic observations generally falling within the corresponding posterior predictive distributions (Fig. S3). These results support the internal consistency of the Bayesian estimation workflow under the assumed model and observation structure. The synthetic data recovery test was conducted using broad priors throughout and without external biochemical constraints. This was intentional, as the purpose of the synthetic test was to assess whether the inference workflow could recover known parameter values under the assumed model structure and observation scheme, rather than to impose additional biological information.

**Figure S3:**
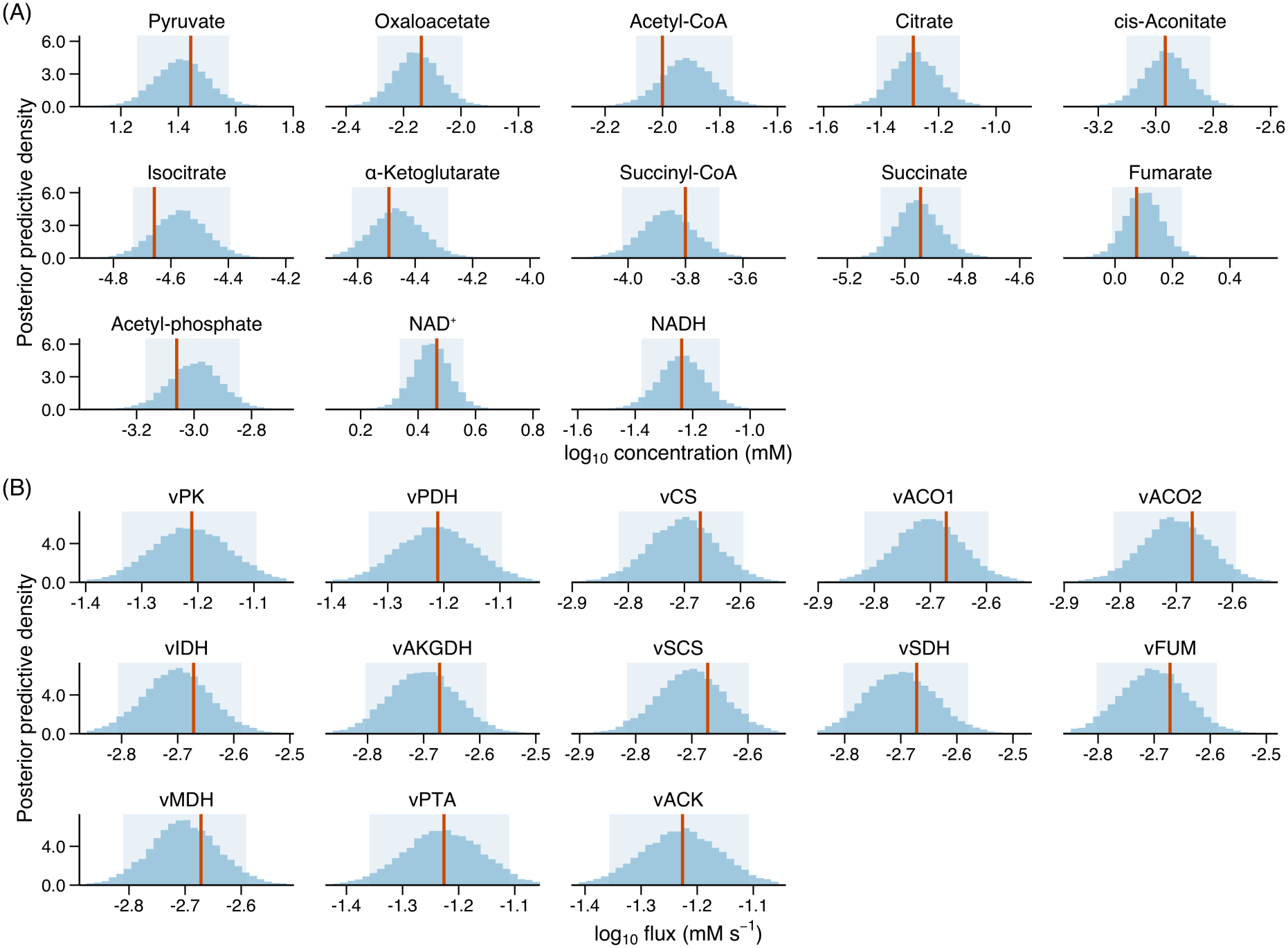
Posterior predictive checks for metabolite concentrations (A) and reaction fluxes (B) in the TCA cycle–fermentation model. Light blue histograms show posterior predictive draws, pale blue shaded regions indicate the central 90% posterior predictive interval, and orange vertical lines denote the synthetic steady-state observation used for model fitting. The comparison assesses whether the fitted model generates metabolite concentrations and fluxes consistent with the synthetic data.

### S2.4 Real-data inference and predictive assessment

Following validation of the inference procedure using synthetic data, the model was calibrated to the experimental steady-state metabolite concentrations and reaction fluxes described in Section S2.1. The computational reliability of the posterior sampling and the predictive adequacy of the calibrated model were assessed in two ways. First, Markov chain Monte Carlo (MCMC) convergence and sampling efficiency were evaluated using chain- and parameter-level diagnostics. Second, prior and posterior predictive distributions were compared to assess how calibration shifted or reduced uncertainty in the model predictions and whether the posterior predictive ensemble was consistent with the observed data.

#### S2.4.1 Experimental-data profile-likelihood analysis and prior specification

Practical identifiability under the actual calibration likelihood was assessed using the experimental steady-state metabolite-concentration and reaction-flux observations. In Figure S4, the profiles for *E*_PK_, *E*_PDH_, *E*_CS_, *E*_ACO_, *E*_IDH_, *E*_AKGDH_, *E*_SDH_, and *E*_ACKA_ crossed the 95% likelihood-ratio threshold on both sides of their maxima. The profiles for *E*_MDH_, *k*, and *α* crossed the threshold on only one side, whereas those for *E*_SCS_, *E*_FUM_, and *E*_PTA_ did not cross the threshold in either direction.

**Figure S4:**
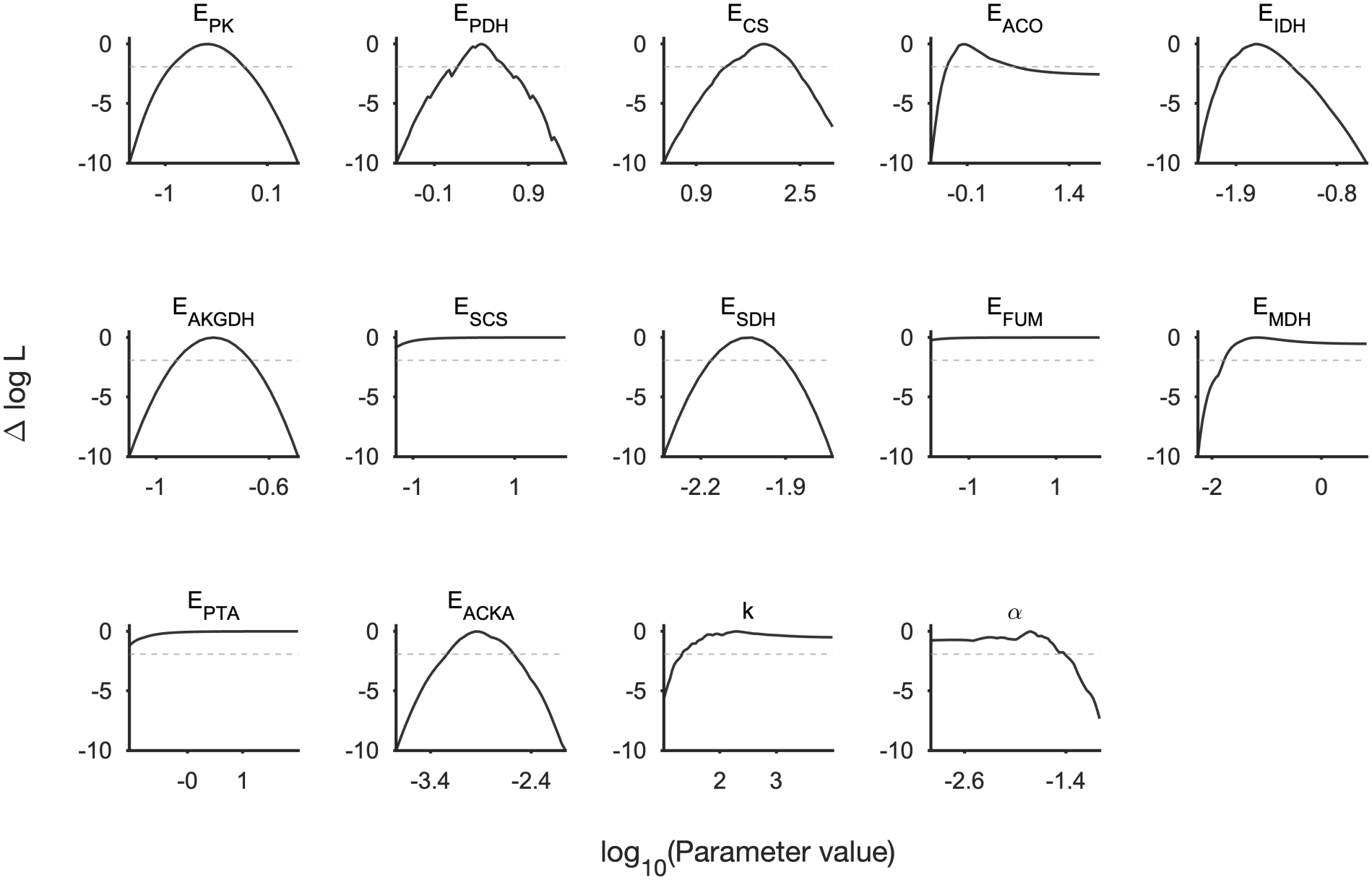
Normalised profile log-likelihoods for the 14 parameters retained after structural identifiability analysis, calculated using the experimental steady-state observations used for Bayesian calibration. Each profile is shown relative to its maximum, such that Δℓ_PL_ = 0 at the maximum-likelihood estimate. The horizontal dashed line denotes the approximate 95% likelihood-ratio threshold, Δℓ_PL_ = *−*1.92.

The experimental-data profile-likelihood classifications and corresponding prior specifications are summarised in Table 3. All 14 structurally identifiable parameters were retained for Bayesian calibration; the profile-likelihood classification affected their prior regularisation rather than their inclusion in the calibrated parameter set.

**Table 3:** Prior distributions used for model calibration. Each prior is specified on the log_10_ scale; the final column gives the corresponding centre on the original scale.

| Parameter | Prior on log<sub>10</sub> scale | Original-scale centre |
|---|---|---|
| *E*<sub>PDH</sub> | 𝒩(−2, 1) | 0.01 |
| *E*<sub>ACO</sub> | 𝒩(−2, 1) | 0.01 |
| *E*<sub>AKGDH</sub> | 𝒩(−2, 1) | 0.01 |
| *E*<sub>SCS</sub><sup>a</sup> | 𝒩(log<sub>10</sub>(0.0062), 0.5) | 0.0062 |
| *E*<sub>SDH</sub> | 𝒩(−2, 1) | 0.01 |
| *E*<sub>PK</sub> | 𝒩(−2, 1) | 0.01 |
| *E*<sub>CS</sub> | 𝒩(−2, 1) | 0.01 |
| *E*<sub>IDH</sub> | 𝒩(−2, 1) | 0.01 |
| *E*<sub>MDH</sub><sup>a</sup> | 𝒩(log<sub>10</sub>(0.0236), 0.5) | 0.0236 |
| *E*<sub>FUM</sub><sup>a</sup> | 𝒩(log<sub>10</sub>(0.0013), 0.5) | 0.0013 |
| *E*<sub>ACKA</sub> | 𝒩(−2, 1) | 0.01 |
| *E*<sub>PTA</sub><sup>a</sup> | 𝒩(log<sub>10</sub>(0.00089), 0.5) | 0.0009 |
| *k* | 𝒩(log<sub>10</sub>(100), 0.5) | 100 |
| *α* | 𝒩(log<sub>10</sub>(0.0195), 0.5) | 0.0195 |
| *σ*<sub>*c*</sub> | 𝒩(log<sub>10</sub>(0.2), 0.15) | 0.2 |
| *σ*<sub>*f*</sub> | 𝒩(log<sub>10</sub>(0.2), 0.15) | 0.2 |

<sup>a</sup>Prior constrained using data from Schmidt et al. [45].

#### S2.4.2 MCMC convergence and sampling diagnostics

Bayesian inference was performed using four independent chains, each with 1,000 warm-up iterations followed by 8,000 sampling iterations, yielding 32,000 post-warm-up draws in total. The No-U-Turn Sampler was run with *adapt delta* set to 0.9 and a maximum tree depth of 12. Convergence and sampling efficiency were assessed using rank-normalised *R* values, bulk and tail effective sample sizes (ESS), divergent transitions, and maximum tree-depth warnings.

For the parameters listed in Table 4, all rank-normalised *R* values were below 1.01, with a maximum of 1.0006. Bulk ESS ranged from 6,194 to 21,752, and tail ESS ranged from 4,327 to 19,098. The lowest bulk and tail ESS values were both observed for *E*_AKGDH_. No divergent transitions occurred among the 32,000 post-warm-up draws. Together, these diagnostics indicate good between-chain convergence and effective sampling for the reported parameters.

**Table 4:** Parameter-level MCMC convergence and sampling diagnostics. Diagnostics were calculated from 32,000 post-warm-up draws obtained from four independent chains, with 8,000 draws retained per chain. Bulk and tail effective sample sizes quantify sampling efficiency in the central and tail regions of each marginal posterior distribution, respectively.

| Parameter | $\widehat{R}$ | Bulk ESS | Tail ESS |
|---|---|---|---|
| *E*<sub>PK</sub> | 1.0005 | 20288 | 16980 |
| *E*<sub>PDH</sub> | 1.0006 | 7114 | 5415 |
| *E*<sub>CS</sub> | 1.0001 | 14757 | 14000 |
| *E*<sub>ACO</sub> | 1.0002 | 14790 | 11521 |
| *E*<sub>IDH</sub> | 1.0003 | 15129 | 12106 |
| *E*<sub>AKGDH</sub> | 1.0005 | 6194 | 4327 |
| *E*<sub>SCS</sub> | 1.0003 | 12445 | 6190 |
| *E*<sub>SDH</sub> | 1.0003 | 14238 | 10728 |
| *E*<sub>FUM</sub> | 1.0002 | 21654 | 16814 |
| *E*<sub>MDH</sub> | 1.0001 | 21752 | 17656 |
| *E*<sub>PTA</sub> | 1.0002 | 10798 | 12102 |
| *E*<sub>ACKA</sub> | 1.0002 | 15742 | 13434 |
| *α* | 1.0002 | 14314 | 17141 |
| *k* | 1.0000 | 17983 | 19098 |
| *σ*<sub>*c*</sub> | 1.0002 | 14343 | 17740 |
| *σ*<sub>*f*</sub> | 1.0003 | 14957 | 14532 |

#### S2.4.3 Prior and posterior predictive checks

Prior and posterior predictive checks were performed to assess the extent to which the observed data updated model predictions (Figure S5). Prior predictive distributions were generated by sampling parameters from their specified priors and propagating these samples through the model; posterior predictive distributions were generated analogously from the post-warm-up posterior draws. Because the same steady-state observations were used for calibration, these checks assess internal consistency rather than out-of-sample predictive performance.

For most metabolic intermediates, the posterior predictive distributions were substantially narrower than their prior counterparts and shifted towards the observed values, indicating that calibration reduced uncertainty in these model predictions (Figure S5A). Succinate showed the largest update, with the posterior predictive distribution displaced by approximately four orders of magnitude relative to the prior. This shift indicates that propagation of the parameter priors through the nonlinear steady-state model placed much of the prior predictive mass below the observed scale, whereas the likelihood strongly constrained the posterior prediction.

The posterior predictive distributions of the reaction fluxes were substantially narrower than their prior counterparts (Figure S5B). Under steady state in this model, the net rates of the nine sequential TCA-cycle reactions constitute a common cycle throughput, while the Pta and AckA rates are equal along the fermentation pathway represented. Consequently, the reaction-specific predictions are strongly coupled by the steady-state mass-balance constraints, and the observed posterior concentration reflects the consistency of the calibrated pathway throughput across the modelled reactions.

**Figure S5:**
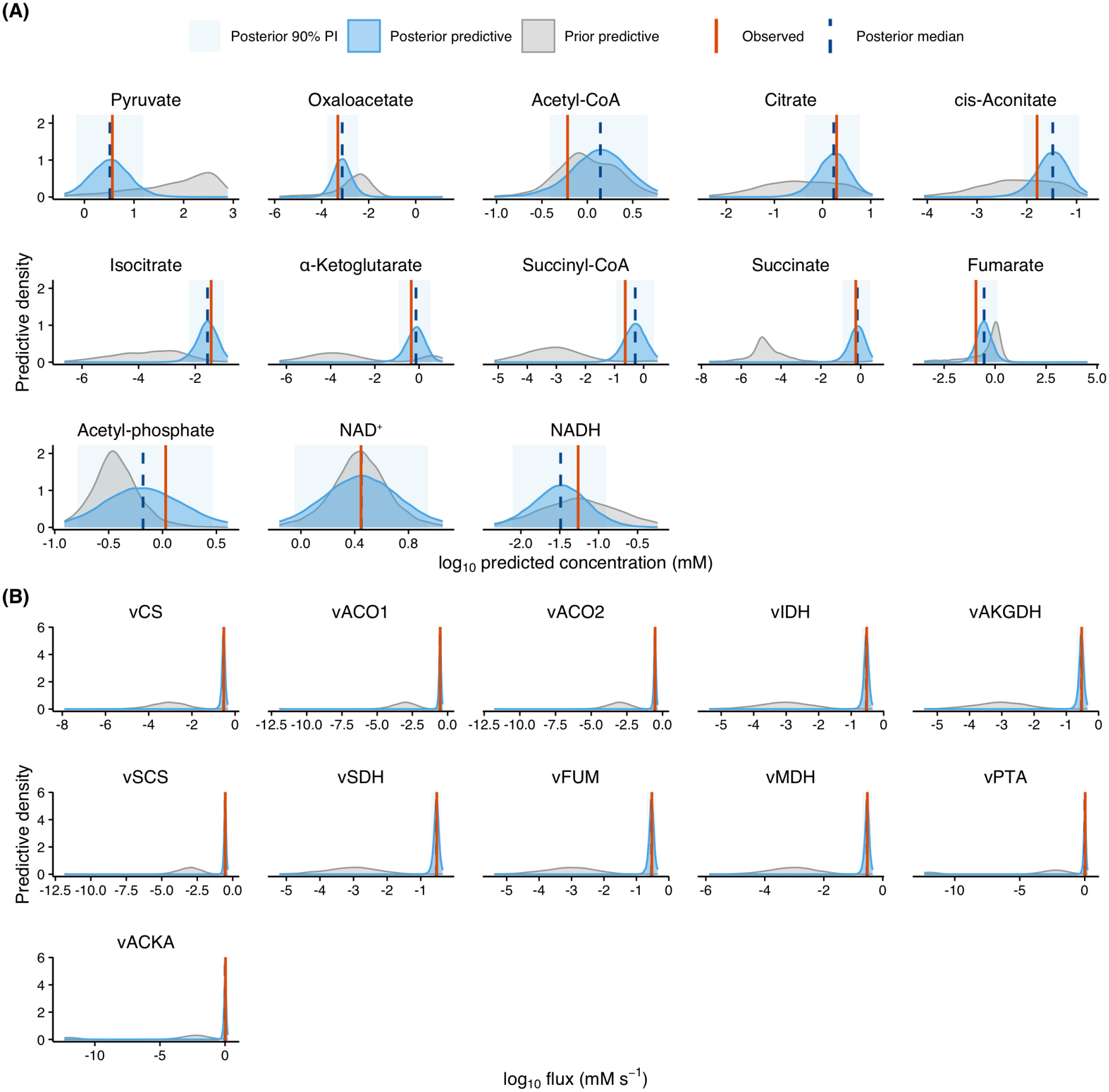
Prior and posterior predictive distributions for the experimental steady-state observations. Distributions are shown for (A) intracellular metabolite concentrations and (B) TCA-cycle and Pta–AckA reaction fluxes. Grey densities represent prior predictive distributions, and blue densities represent posterior predictive distributions obtained after calibration to the experimental data. Dark-blue dashed vertical lines indicate posterior predictive medians, pale-blue shaded regions indicate central 90% posterior predictive intervals, and orange solid vertical lines indicate the observed values. Concentrations and fluxes are shown on the (log_10_) scale in mM and (mMs^*−*1^), respectively. The comparison shows how calibration shifts and narrows the model predictions relative to their prior distributions.

